# Supervised white matter bundle segmentation in glioma patients with transfer learning

**DOI:** 10.1101/2025.06.13.659452

**Authors:** Chiara Riccardi, Ludovico Coletta, Sofia Ghezzi, Gabriele Amorosino, Luca Zigiotto, Jorge Jovicich, Silvio Sarubbo, Paolo Avesani

## Abstract

In clinical settings, the virtual dissection of white matter tracts represents an informative source of information for monitoring neurological conditions or to support the planning of a treatment. The automation of this task using data-driven approaches, and in particular deep learning models, provides promising evidence of good accuracy when carried out on healthy individuals. However, the lack of large clinical datasets and the profound differences between healthy and clinical populations hinder the translation of these results to patients. Here, we investigated for the first time the effectiveness of transfer learning in adapting a deep learning architecture trained on a healthy population to glioma patients. Importantly, we provided the first thorough characterization of domain shift and its complexity, distinguishing systematic (i.e. measurement and pre-processing related) from tumor specific components. Our results suggest that (i) models trained on a large normative healthy population have a significant performance drop when the inference is carried out on patients; (ii) transfer learning can be an effective strategy to overcome the shortage of clinical data and to manage the systematic shift; (iii) fine tuning of the learning model cannot accommodate large white matter deformations induced by the tumor. The results were coherent across the five white matter bundles and three different input modalities tested, highlighting their robustness and generalizability. Our work provides valuable insights for advancing automated white matter segmentation in clinical populations and enhancing clinical transfer learning applications.

**Graphical Abstract:** 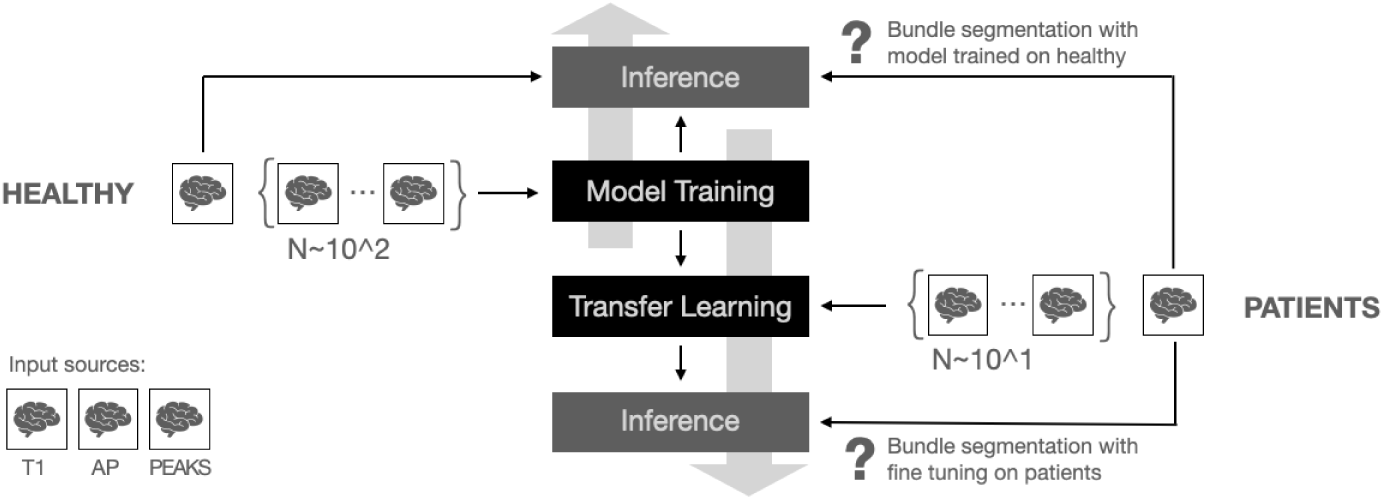

**Highlights:**

- White matter bundle segmentation is essential for planning glioma tumor resection.
- We trained a deep learning model on a healthy dataset to segment white matter bundles.
- We observed a sub-optimal performance on glioma patients due to domain shift.
- Transfer learning mitigated domain shift between healthy and clinical populations.
- We highlighted domain shift domains solved/not solved by transfer learning.

## 1. Introduction

Magnetic resonance imaging (MRI) has emerged as a valuable non-invasive tool to characterize the large-scale architecture of the brain. Structural MRI provides insights into brain’s morphology, while diffusion weighted imaging (DWI) allows to track white matter pathways in-vivo (Wang and He, 2024; Le Bihan, 2003).

Of note, single white matter pathways do not exist in isolation, but group into coherent formations known as bundles. Their identification is of pivotal importance, as white matter bundles (or tracts) support the characterization of healthy functioning as well as the derangement occurring in neurological disorders (Yamada et al., 2009; Kamagata et al., 2021; Ghazi et al., 2023). For brain tumor patients, the segmentation of white matter bundles - also known as virtual dissection - is particularly important, as it provides neurosurgeons with a neuro-biologically grounded reference driving the selection of brain areas not to be damaged during surgery (Yamada et al., 2009; Ghazi et al., 2023). As such, virtual dissection plays a crucial role in avoiding surgery-induced permanent cognitive impairments (Yang et al., 2021).

White matter bundle segmentation is typically performed manually using the patient’ specific MRI acquisition, combining the identification of anatomical landmarks and the manual selection of white fibers reconstructed via DWI and fiber tracking (Andica et al., 2023; Wakana et al., 2007). In the neurosurgical theater, the virtually dissected bundle is then used as an anatomical reference driving tumor resection during surgery (Cho et al., 2014). Given the requested amount of knowledge about human brain anatomy and the dramatic consequences arising from an erroneous segmentation, the pre-surgical identification of clinically relevant white matter tracts is the responsibility of dedicated and ad-hoc trained personnel. Nonetheless, manual bundle segmentation is time-consuming, prone to errors, and highly operator dependent (Tunç et al., 2016; Joshi et al., 2024).

To address these critical issues, the research community has produced a significant effort to develop automated and reliable methods for the bundles dissection (Yendiki et al., 2011; Ghazi et al., 2023).

Automated methods for white matter bundles segmentation can be classified into two main strategies: atlas-based and learning-based. The former exploits manually curated and normative references delineating prototypical bundle representations (Garyfallidis et al., 2018; Zhang et al., 2018). Learning-based models, on the other hand, use a collection of ground-truth examples to automatically extract the set of rules governing the segmentation process (Wasserthal et al., 2018; Bertò et al., 2021). In the context of glioma research, atlas based methods perform poorly, as they cannot account - by definition - for the deformation induced by the presence of a brain tumor. Analogously, the development of learning based methods for white matter tracts segmentation in glioma patients is hindered by the lack of dedicated large-scale and publicly available clinical datasets. Taken together, a reliable solution to automatically extract clinically relevant white matter bundles in brain tumor patients is currently missing but it is an urgent clinical needed.

A conceptually simple strategy to address this gap is learning a model on a reference population of healthy individuals, and then adapting to a custom (and usually much smaller) sample of interest. This strategy - known as transfer learning - yielded promising results across several clinical neuro-imaging domains, as it allows to compensate for a difference in data distribution across samples - the so-called domain shift (Ismaila et al., 2022). However, transfer learning has never been investigated in the context of white matter bundles segmentation in glioma patients.

Leveraging the unprecedented availability of manually curated tracts segmentation in glioma patients, in the present work we investigated the efficacy of transfer learning in the context white matter bundle segmentation in this clinical sample. Specifically, we first trained a convolution neural network on a large-scale dataset encompassing hundreds of healthy individual and several manually annotated white matter bundles. We then applied transfer learning to set of glioma patients. The main contributions of our work are twofold: (i) We developed a novel metric enabling the distinction and quantification of domain shift occurring due to the tumor (biological bias) from systematic variations (different MRI scanners, acquisition parameters, and pre-processing, systematic bias) with single subject resolution. (ii) Importantly, we show that transfer learning could recover the domain shift related to systematic variations, but not the spatial re-arrangement of the bundles induced by the tumor.

Our findings were coherent across the five white matter bundles and three different input modalities tested, highlighting their robustness and generalizability.

Taken together, our results advance the state-of-the-art by showing that white matter bundle segmentation in glioma patients is a challenging task that cannot be addressed with simple transfer learning approaches.

## 2. Related work

With respect to glioma patients, the development of transfer learning solutions for automated bundles dissection has been hampered by the specific unavailability of manually curated segmentations, resulting in the direct application of solutions originally developed for healthy individuals. (Richards et al., 2021; Garyfallidis et al., 2018; Zhang et al., 2018, 2020; Xue et al., 2023; Bertò et al., 2021). Notably, one study tried to develop a glioma-to-glioma learning procedure (Moshe et al., 2022). Although promising, the study reported lower accuracy in the segmentation of the pyramidal tract compared to our results obtained with the transfer learning method we propose in the present work.

While transfer learning has been poorly investigated in the context of bundles segmentation for glioma patients, some notable applications in healthy individuals have been recently reported. In(Lu et al., 2022; Liu et al., 2023), the authors explored the effectiveness of fine-tuning and data augmentation strategies in developing a solution able to generalize across anatomically different bundles and when few target examples are available, highlighting the potential of transfer learning.

However, none of the reports considered in this overview investigated the differential impact of neurobiological factors (such as tumour localization) and more systematic variations (different MRI scanners, acquisition parameters, pre-processing, and segmentation procedure) in determining the efficacy of the transfer learning task. In the present report, we fill this gap by introducing a conceptually and methodological simple set of metrics that can be used at the single subject level, providing the first investigation of a transfer learning strategy for automated bundles dissection in glioma patients.

## 3. Materials

### 3.1. The Datasets

We considered two datasets in this study: a public dataset of healthy subjects, and a dataset of glioma patients.

### 3.2. The healthy subjects’ dataset

Healthy subjects data were obtained from TractoInferno^1^, a large open access collection of hundreds of MRI acquisitions, deriving from 6 sites. The dataset includes T1-weighted (T1w), diffusion-weighted imaging (DWI), and their derivatives (Poulin et al., 2022). All TractoInferno scans were acquired using 3T scanners, with acquisition parameters such as b-value, resolution, TE, TR, and the number of gradient directions varying between the 6 acquisition sites, as described in detail in (Poulin et al., 2022). The data were distributed already preprocessed. For DWI images, preprocessing included brain extraction, denoising, artifact correction for eddy current and N4 bias field, normalization, and resampling; for T1w images, preprocessing included denoising, N4 bias field correction, and registration. Rigorous multi-step manual quality control was performed to remove poor quality and artifactual images (Poulin et al., 2022).

For each subject, Poulin et al. (2022) fitted a fiber orientation distribution function (fODF) to the DWI data using constrained spherical deconvolution (Tournier et al., 2007), from which an ensemble tractography was derived. The ensemble tractography was obtained by merging four tracking techniques: deterministic (Basser et al., 2000), probabilistic (Tournier et al., 2012), particle filtered (Girard et al., 2014), and surface-enhanced tracking (St-Onge et al., 2018). 30 White matter bundles were automatically segmented using RecoBundlesX (Garyfallidis et al., 2018; St-Onge, 2021), and then manually checked visually. In case of inaccuracies in the anatomical representation of a bundle, the authors removed it from the dataset. As a result, not all subjects have all 30 segmented fiber bundles (Poulin et al., 2022).

### 3.3. The Clinical dataset

The clinical dataset comprises MRI data of glioma patients, collected at the Department of Neurosurgery of the University Hospital S. Chiara in Trento. We considered a sample of 21 patients with a brain tumor in the left hemisphere. Each patient underwent an MRI examination as part of the standard protocol for glioma surgery. T1 and DWI sequences were acquired for preoperative planning of glioma resection using a GE 1.5T clinical scanner. T1-weighted volumetric sequences were acquired with the following parameters: axial acquisition, square field of view (FOV) of 256 mm, FA = 12°, voxel size=1×1×1*mm*, TR/TI/TE = 10.64/450/4.23 ms (Zigiotto et al., 2022). The DWI images were acquired in a single acquisition with 60 directions, using a single-shot multislice spin-echo-echo-planar sequence with the parameters: 50 slices; square FOV 240 mm; voxel size of 2.4×2.4×2.4 *mm*; TR/TE = 13000/95.8 ms; b values of 0 and 1,000 *s*/*mm*^2^; FA of 90° (Zigiotto et al., 2022).

Using anatomical images as a guide, tumor volume masks were manually segmented and validated by neuroradiology experts [L.Z] using MRIcroGL software (Rorden and Brett, 2000; Rorden and Halchenko, 2024). DWI data were processed through a multi-step pipeline (Zigiotto et al., 2024). First, DWI data were preprocessed using the Tortoise toolkit (Irfanoglu et al., 2017), to correct for artifacts such as DWI thermal noise, Gibbs ringing, eddy current, motion distortion, and susceptibility-induced EPI distortion. Then, the voxel-wise white matter fiber orientation distribution function (fODF) was estimated using the constrained spherical deconvolution (CSD) technique (Tournier et al., 2007) to perform probabilistic tractography (Tournier et al., 2012). White matter fiber bundle segmentation was performed by an expert operator [L.Z] using Trackvis software^2^. In this process, an expert neurosurgeon [S.S] manually defined multiple regions of interest (ROIs) based on anatomical regions identified by using T1-weighted MRI as a reference image and supported by the color-coded FA map (Mori et al., 2002; Catani and Thiebaut de Schotten, 2008). Fiber bundles were then manually refined using the Tractome interactive tool (Porro-Muñoz et al., 2015).

### 3.4. The white matter bundles

In our study, we considered 5 fiber bundles available in both the clinical and the TractoInferno datasets. Specifically, these bundles were located in the left hemisphere: the pyramidal tract (PYT_L), the arcuate fasciculus (AF_L), the frontal aslant tract (FAT_L), the inferior fronto-occipital fasciculus (IFOF_L), and the inferior longitudinal fasciculus (ILF_L). While these bundles were consistently segmented in all subjects in the clinical dataset, not all subjects in TractoInferno had all of these bundles due to the aforementioned exclusions. The detailed distribution of these bundles in both datasets is shown in Table 1.

**Table 1:**
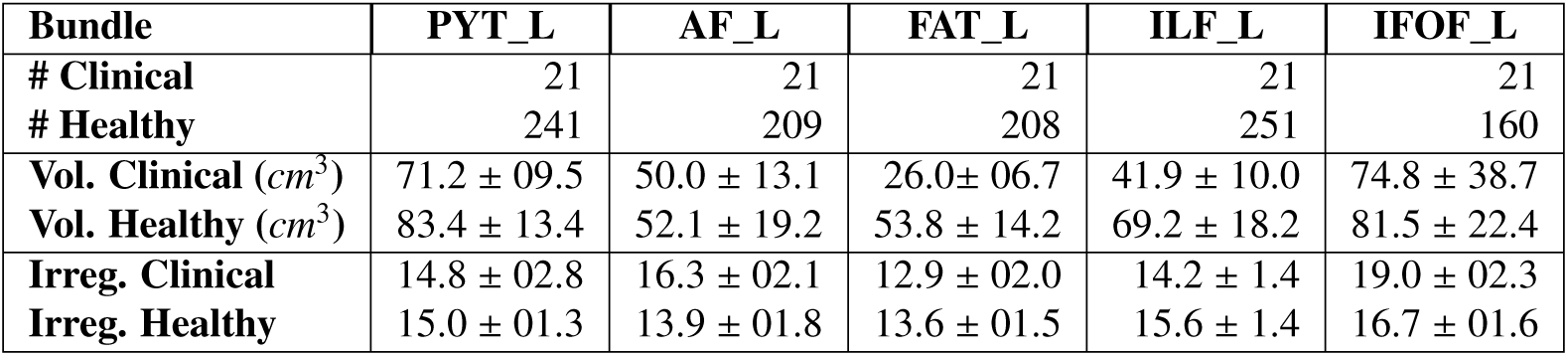
White matter bundle statistics for the Healthy and Clinical datasets (denoted respectively by *Heathy* and *Clinical*). The statistics include: the number of available white matter bundle segmentations (rows: *#Clinical* and *#Healthy*); the mean and standard deviation of bundle volumes in *cm*^3^ ( rows: *Vol. Clinical (cm*^3^) and *Vol. Healthy(cm*^3^)); the mean and standard deviation of the Irregularity as defined by Yeh (2020) (rows: *Irreg. Clinical* and *Irreg. Healthy*). The columns indicate the white matter bundles: the pyramidal tract (*PYT_L*), the arcuate fasciculus (*AF_L*), the frontal aslant tract (*FAT_L*), the inferior fronto-occipital fasciculus (*IFOF_L*), and the inferior longitudinal fasciculus (*ILF_L*).

## 4. Methods

### 4.1. The Transfer learning method

In the following paragraphs, we will introduce how the transfer learning was applied in our work to foster bundle segmentation in glioma patients.

We adopt the same definitions of transfer learning, domain, and task used by Pan and Yang (2010). The domain is defined as 𝒟 = {*X*, *P*(*X*)} (Pan and Yang, 2010). *X* is the set of the possible input values. In the case of bundle segmentation, *X* includes the possible values of the voxels, in the images used as input for the segmentation models. *P*(*X*) is the probability distribution of these values.

The task 𝒯 is defined as 𝒯 = {*Y*, *f* (·)} (Pan and Yang, 2010). *Y* is the label space. In the specific case of bundle segmentation, *Y* includes the values 1 and 0, indicating for each voxel whether the bundle is respectively present or not. *f* (·) is the objective function that should be learnt from data and consists of pairs of *x_i_*, *y_i_*, with *x_i_* ∈ *X* and *y_i_* ∈ *Y*. This function predicts the corresponding *y_i_*, by taking as input each *x_i_*, and can also be denoted in probabilistic form as *P*(*Y*|*X*) (Pan and Yang, 2010).

In transfer learning, a source and target domain are commonly considered, namely 𝒟*_S_* and 𝒟*_T_*. For each of them, the *data* sampled from the respective domain and used to learn the objective function can be considered.

The source domain data is defined as 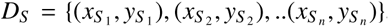, with 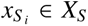 and 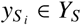. The target domain data is defined similarly, as 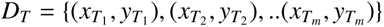, with 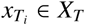 and 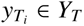 (Pan and Yang, 2010). Often the target domain data has fewer examples than the source domain data.

The goal of transfer learning is to improve the objective function *f_T_* (·) in the target domain 𝒟*_T_*, using the knowledge of a source domain and task, 𝒟*_S_* and 𝒯*_S_*, and assuming that 𝒟*_S_* * 𝒟*_T_*, or 𝒯*_S_* * 𝒯*_T_* (Pan and Yang, 2010).

#### 4.1.1. The source and the target tasks

The source task and the domain that we consider (*T_S_* and *D_S_* ) refer to the segmentation of the bundles in the healthy subjects’ dataset. The target task and domain that we consider (*T_T_* and *D_T_* ) refer to the segmentation of bundles in the glioma patients’ dataset. In this work, we test the hypothesis that transfer learning improves the learning of the target task, *T_T_* the bundle segmentation in the patients, by exploiting the knowledge of the source task, *T_S_* the bundle segmentation in the healthy population.

#### 4.1.2. The learning models

Although numerous automated models for white matter bundle segmentation have been documented in the literature (Yendiki et al., 2011; Wasserthal et al., 2018; Garyfallidis et al., 2018; Bertò et al., 2021), we focused our analysis on methods utilizing volumetric representation. This decision was made to minimize the variance in bundle representation deriving from the different tractography algorithms.

As a reference method for the volumetric representation of bundle segmentation, we chose a 3D U-Net deep-learning model (Isensee et al., 2021).

While we used the same architecture for the convolutional neural network for our analysis, the 3D U-Net, we designed three different training strategies.

The first strategy was the standard training process with randomly initialized model weights(*M-train*). This strategy was used to train the bundle segmentation model on the large healthy dataset.

The second and third training strategies adapted the model resulting from *M-train* to a new set of data with fine-tuning (in our case the glioma patients dataset). These are referred to as *M-retrain* and *M-transfer*.

In *M-retrain*, or retraining, the network weights were initialized using the final epoch of *M-train* and then were all further optimized with new training data. This could be considered as the baseline strategy for handling the domain shift when moving from a healthy to a clinical setting. *M-transfer*, or transfer learning (Sundaresan et al., 2021), was similar to the retraining strategy but used a different weight update schema. In this approach, learning was limited to the decoder of the 3D U-Net, which reconstructs the original space from the latent space (Sundaresan et al., 2021).

We treated each bundle and input modality separately in the transfer learning and retraining process. For example, the model optimized for segmenting the pyramidal tract (PYT_L) in healthy subjects using T1w input served as the source to transfer knowledge to improve the model optimized for segmenting the same bundle (PYT_L), with the same input modality (T1w) in patients. This approach was applied similarly to other bundles and input modalities.

#### 4.1.3. The data modalities

We trained our models using three different and increasingly complex inputs: T1w images, anisotropic power, and the three main peaks of the fiber orientation distribution function (fODF). The motivation was to investigate if and how the transfer learning is influenced by the use of different input image modalities. The selected input modalities have already been used as source data for the bundle segmentation. T1w input images have been extensively tested in a recent work (Yang et al., 2022), where the authors provided robust evidence that they can be a reliable source to estimate the spatial distribution of the bundles and a viable alternative when DWI data are not available, such as in clinics. The three main fODF peaks extracted from CSD are the input data used by the popular method TractSeg (Wasserthal et al., 2018). Anisotropic power (Dell’Acqua et al., 2014) can be informative of the fiber pathways. Pomiecko et al. (Pomiecko et al., 2019) showed that using anisotropic maps as input of a Convolutional Neural Network (CNN) is effective for segmenting white matter bundles.

### 4.2. Performance evaluation

We used two metrics to measure the performance of the learning models optimized with the different training strategies. The first metric was the Dice Score Coefficient (DSC), which assessed the voxel-wise similarity between the predicted bundle mask and the corresponding manually annotated ground truth.

The second metric is referred to as the *Improvement Map*. This metric pairwise compared two training strategies at the voxel level when performing inference on the same dataset to evaluate spatially where one training strategy outperformed the other.

To compute the Improvement Map, we analyzed a set of brain images of different individuals, each linearly registered to MNI. Two models took each the same brain image as input and generated two bundle masks as bundle segmentation predictions for the same individual. Each predicted segmentation was then converted into a binary true prediction map. This map had a value of 1 for voxels with either positive or negative true predictions, and 0 otherwise.

We compared the true prediction maps generated by the two competing models to understand where one model outperformed the other. In particular, we defined an individual Improvement Map as the difference between the true prediction maps of the two models for the same image. If both models performed identically, the voxels of the Improvement Map had a value of 0; otherwise, the voxels had a value of 1 or −1, indicating a difference in performance.

By averaging the individual Improvement Maps voxel-wise across the population, we obtained a global Improvement Map. Considering two models, positive values in the global Improvement Map indicated better performance by the first model, while negative values indicated better performance by the second model. Values near to 0 indicated similar behavior. The Improvement maps are shown in Figure 5.

### 4.3. The Domain shift

#### 4.3.1. The categories of shift

In our study, we measured the shift between the data collected in the source domain and the data of the target domain, referred to as *D_S_* and *D_T_*, defined in Section 4.1. The considered source domain dataset was the healthy population dataset, while the target domain was the glioma patient sample. For simplicity, we refer to this data domain shift as *Domain shift*.

We categorized domain shift into *tumor-related shift* and *systematic shift*. In doing so, we tried to capture their differential impact on the data: 1)*tumor-related shift* is highly inhomogeneous because tumors have different locations and volumes across patients, as shown in Figure 1. Thus, the tumor causes anatomical deviations from the normative model that are highly subject-specific. 2) the *systematic shift* is caused by differences that are constant and homogeneous across the two datasets. This domain shift is caused by the different scans (3T versus 1.5T), the different acquisition parameters, the different bundle annotation protocols (manual ROI-based versus RecobundleX with visual checking), and the different tractography algorithms used (probabilistic versus ensemble tractography).

**Figure 1:**
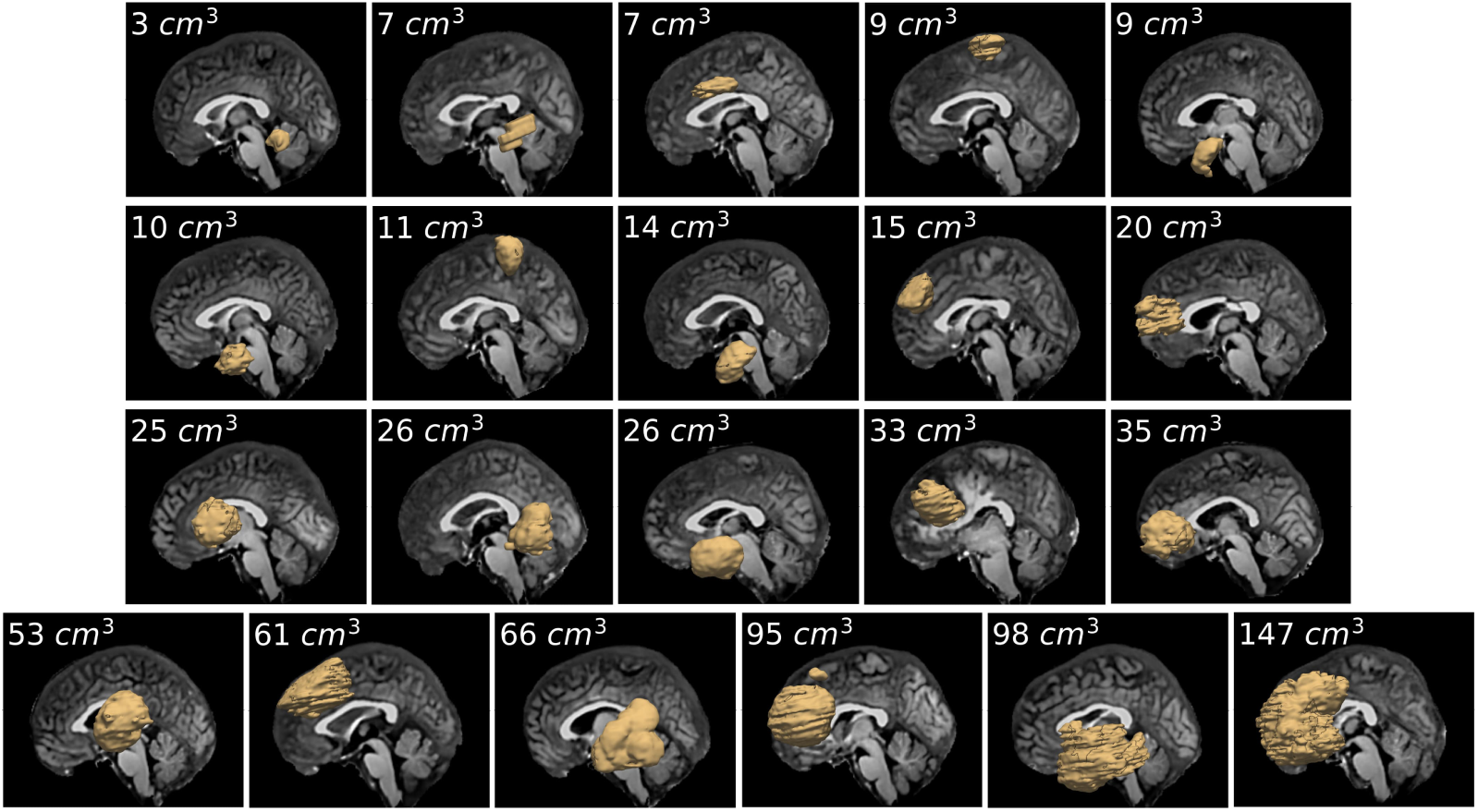
Clinical dataset. Visualization of each patient’s tumor mask overlaid on the T1-weighted image, in the left hemisphere, with the corresponding tumor volume reported in cubic centimeters *cm*^3^

#### 4.3.2. Domain shift assessment

We assessed the domain shift between the datasets of the healthy population and of the glioma patients using two approaches. In the former approach, the spatial distribution of the bundle masks was measured using probabilistic maps. For each bundle, two probability maps of the bundle masks were computed, one for the healthy individuals, and the other for the glioma patients. Then, the absolute difference between the two probabilistic maps was computed, accounting for the shift between the two data samples for each bundle.

The second metric, based on tractography, aimed to highlight the shift of the bundle trajectories. In our study, the trajectory of a bundle was encoded as the most representative fiber of the whole bundle and is referred to as the *skeleton* (Li and Hu, 2013) of the bundle. The skeleton was computed as the average pathway in the area of maximum fiber density of the bundle (Amorosino et al., 2023). The intuitive idea was to measure the difference between the trajectories of two bundles, along the points of the skeleton, by considering the warp of the registration between the two bundles. For this purpose, we introduce the concepts of *Warp V-Norm* and *Warp V-Norm Profile*.

The *Warp V-Norm* estimates the local shape differences between two bundles, encoded as volumetric binary masks, and it was computed in several steps. First, the two bundles were spatially normalized in a common space using an affine transformation, derived from the registration of the related T1-w images. Then rigid and diffeomorphic transformations were estimated to optimize the matching between the bundles encoded as volumetric binary masks. This transformation encoded local spatial differences since the more a certain bundle’s portion was moved during the registration, the more it differed from the fixed bundle. We measured this local spatial difference with the Warp V-Norm, defined as the 2-norm of the vector encoding the entire transformation in each voxel of the warped image.

The Warp V-Norm is defined in each voxel of the moving bundle warped after the registration and is difficult to visualize. For this reason, we considered the skeleton fiber, estimated from the bundle fixed in the registration. We mapped the Warp V-Norm values along each point of the skeleton, considering the corresponding voxels. We obtained a vector summarizing the Warp V-Norm values along the main trajectory of the bundle. This vector is referred to as the *Warp V-Norm profile*.

We estimated the Warp V-Norm and the Warp V-Norm profile to pairwise compare each subject’s bundle with a normative bundle model. Specifically, we computed the registration between a bundle normative model (fixed image) and: 1) each bundle of the clinical dataset (moving images), 2) each bundle of the test-set of the healthy subjects dataset (moving images). High values of Warp V-Norm and of Warp V-Norm profile indicated a large deviation of a given bundle from the underlying normative model.

## 5. Experiments and Results

### 5.0.1. The Data Preparation

First, we prepared the inputs to train the models, namely the ground truth segmentation of the bundles and each one of the three image data modalities considered for the learning.

The ground truth segmentation used as models’ input was obtained by encoding the fiber representation of each segmented bundle in the corresponding voxel mask, defined in the subject’s T1-weighted image-space voxel grid.

The three data modalities used were: (i) the T1w image (Yang et al., 2022), (ii) the anisotropic power (Pomiecko et al., 2019), and (iii) the three major peaks extracted from the Constrained Spherical Deconvolution (CSD) model (Wasserthal et al., 2018). The peaks of the spherical harmonic function were extracted in each voxel of the CSD models using the MRtrix toolbox (Tournier et al., 2019). The anisotropic power (Dell’Acqua et al., 2014) was calculated using Scilpy ^3^, with the fitting of the Constant Solid Angle (CSA) model (Aganj et al., 2010) on the DWI data. All the data were normalized to MNI space using a linear transformation (ANTs (Avants et al., 2011)). The linear transformation was estimated by performing an affine registration between the individual subject’s T1-weighted image and a T1-weighted template in MNI space (Louis, 2015). Statistics on the number and volume of white matter bundles used in this study are reported in Table 1.

### 5.1. Domain shift characterization

Our first analysis measured the domain shift between the two datasets. We first mapped the tractography representation of the bundles to voxels, obtaining binary masks. Subsequently, we normalized the bundle masks to the MNI space via an affine registration of corresponding anatomical T1w images. We then aggregated all the volumetric representations of each bundle to obtain probability maps measuring their the spatial distribution. We generated two probability maps for each one of the bundles of interest: one for the healthy population and one for the glioma patients. To highlight the difference in the spatial distribution between the two samples of data, we generated an additional map that reports the absolute difference between them, as reported by Figure 2. The visual inspection of this latter map allows to detect consistent differences in the spatial organization of the bundles, as higher values account for differences that are particularly consistent across the entire dataset, and reflect systematic domain shift. The probability maps for each bundle and population, together with the corresponding difference maps are shown in Figure 2.

**Figure 2:**
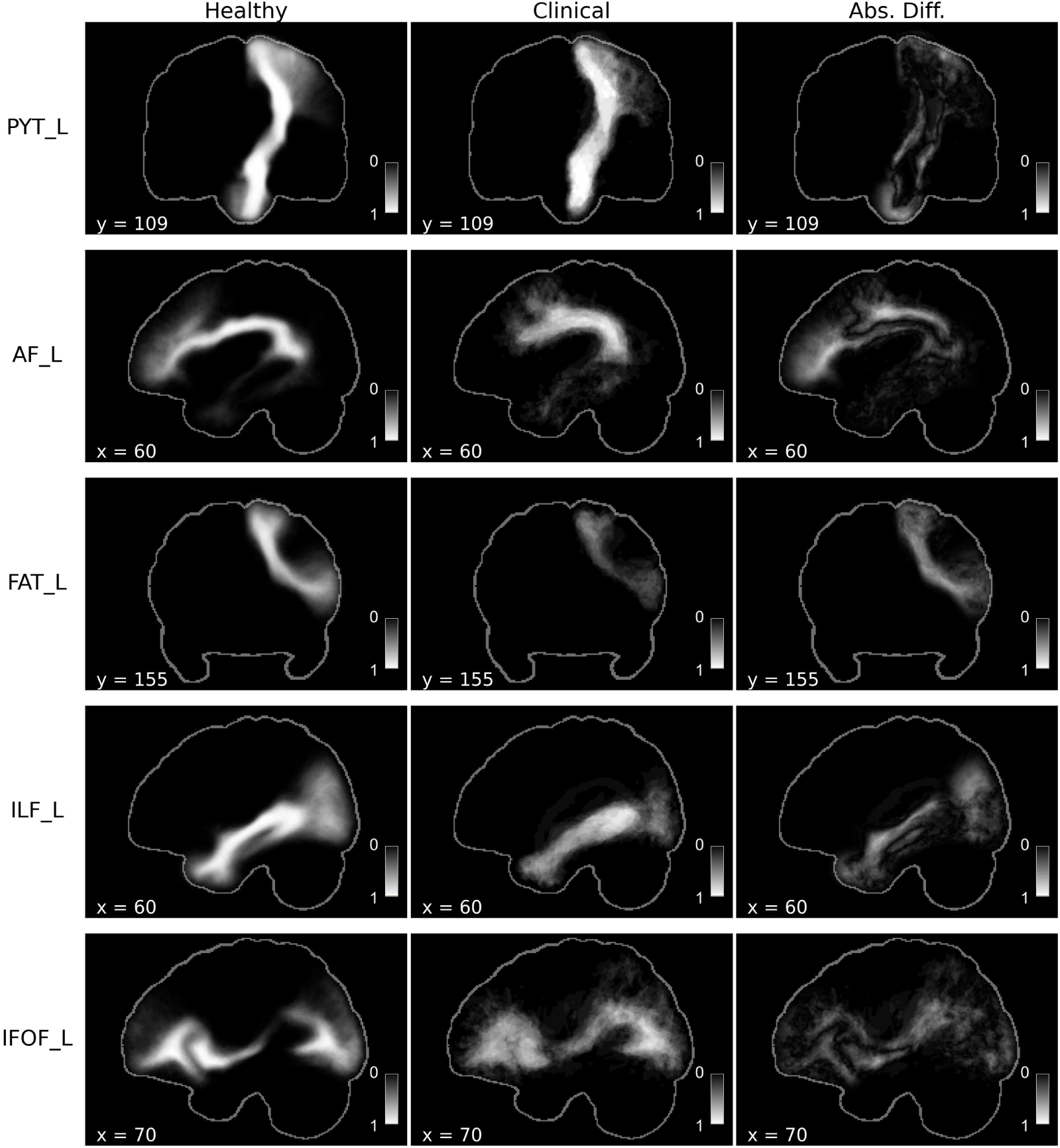
Domain shift. In the 1st column: the probability map for the healthy population (TractoInferno); in the 2nd column: the probability map in the Clinical dataset with glioma patients; in the 3rd column: the absolute difference between the two probability maps. The rows indicate the white matter bundles: the pyramidal tract (*PYT_L*), the arcuate fasciculus (*AF_L*), the frontal aslant tract (*FAT_L*), the inferior longitudinal fasciculus (*ILF_L*), and the inferior fronto-occipital fasciculus (*IFOF_L*).

To measure the domain shift according to the tractography representation, we computed for each bundle the Warp V-Norm profile for individuals of the healthy population in the test sets, and for all the glioma patients. As a reference template of a bundle, we estimate a normative model from the probability maps of the healthy population, thresholded so that the mask volume matches the mean volume of the whole healthy individuals. We use this template as a reference to compute a rigid and a diffeomorphic transformations of each bundle mask using ANTs (Avants et al., 2011). These transformations - mapped to the skeletons of the bundles - provide a measure of the displacement of the main pathway with respect to the normative model, separately for the healthy individuals and the patients, as shown in Figure 6. The Figure highlights the individuals contributing the most to the so-defined domain shift.

### 5.2. The training strategies and inference evaluation

For the implementation of 3D U-Net, we used nnU-Net (Isensee et al., 2021), a state-of-the-art deep learning framework for biomedical image segmentation. To reduce the computational load and balance the classes ratio of the binary classifiers, we resized the bounding box of the images, so that for each bundle we excluded the contralateral hemisphere, and brain regions in the same hemisphere not compliant with anatomical plausibility priors. For each bundle and each input image modality, we trained the model in three separate sessions, obtaining a total of 45 different segmentation models (the combination of 3 training strategies, 3 input images, and 5 bundles). In the first session, a 3D U-Net model was trained with healthy data (M-train); in the second session, the M-train model was retrained with patients’ data only (M-retrain); in the third session, the M-train model was fine-tuned with a transfer learning procedure using patients’ data only (M-transfer). Each of these three models was applied for inference on both healthy and patient test sets. To measure models’ performance, we computed the average DSC between predicted and ground true bundle masks, as reported in Table 2 and Figure 4.

**Table 2:**
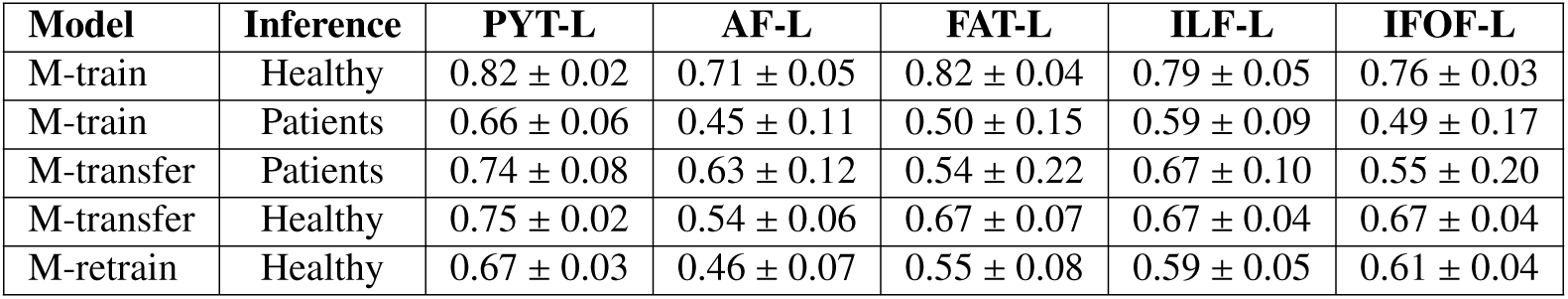
Learning strategies evaluation. Mean and standard deviation of DSC between ground truth and predicted bundle segmentation of models M-train only trained on healthy, or trained on healthy and on glioma patients with M-transfer, or M-retrain by using as input T1 images. We distinguish whether the models are used for inference on healthy individuals or patients. The bundles considered are: the pyramidal tract (*PYT_L*), the arcuate fasciculus (*AF_L*), the frontal aslant tract (*FAT_L*), the inferior longitudinal fasciculus (*ILF_L*), and inferior fronto-occipital fasciculus (*IFOF_L*).

The design of splitting in train and test is different for the healthy individuals and patients data. For the healthy individuals dataset, the split had an 80% − 20% ratio. Since the number of individuals having a certain bundle was not constant in the healthy dataset (see Table 1), the train/test set partition was computed independently for each bundle. A 3-fold cross-validation was performed with 50 epochs each.

Due to the small sample size, Training sessions on patients were designed instead with a stratified cross-validation scheme. Specifically, the data were partitioned according to a 5-fold cross-validation, and a nested 3-fold cross validation was performed with 200 epochs for each training. The variance in the spatial distribution of the bundles was much larger for the patients, and random splitting might have resulted in folds unbalanced for tumor-induced alterations. Therefore, the partitioning design in the 5-folds was revised to balance the distribution of bundles with major alterations. We measured the alteration by calculating the Warp V-Norm profile, as was reported in the Methods section. The nnU-Net framework divided the images into mini-batches (Isensee et al., 2021). We set the number of mini-batches so that each epoch selected each image of the training set only once. The intensity normalization of the input images was embedded in the nnU-Net and performed via a z-scoring.

### 5.3. Input modalities and segmentation evaluation

We compared the segmentation performance across input modalities for each bundle. We first focused our analysis on the baseline situation where the model was trained on the healthy population and the inference was performed on healthy individuals. In particular, we performed a one-sided Wilcoxon signed-rank test to compare the paired distributions of the DSC of predictions on the same test set when the models where trained with different input modalities. The DSC distributions were significantly different for all bundles when comparing AP versus peaks and T1w versus peaks: the models trained on AP and T1w performed better than models trained on peaks (see Supplementary Material for a detailed tests’ summary report). We found that models trained on T1w perform significantly better than models trained on AP, *p*-value < 0.01, for all the bundles except AF_L. The one-sided Wilcoxon signed-rank tests to compare models performances were computed with Scipy ^4^, and the values of the statistics and of the *p*. values are reported in the Supplementary materials in Table S2. The comparison of DSC distributions for the healthy population is shown in Figure 3. This figure shows how the magnitude of the difference was greater for the comparisons of AP versus peaks and T1 versus peaks than for T1 versus AP. We also computed the improvement map to spatially visualize where models with T1 and AP inputs behaved more accurately than those with peaks input, and where T1 input achieves better performance than AP input. A visualization of the Improvements maps, thresholded for values greater than zero, is reported in the supplementary materials in Figure S1.

**Figure 3:**
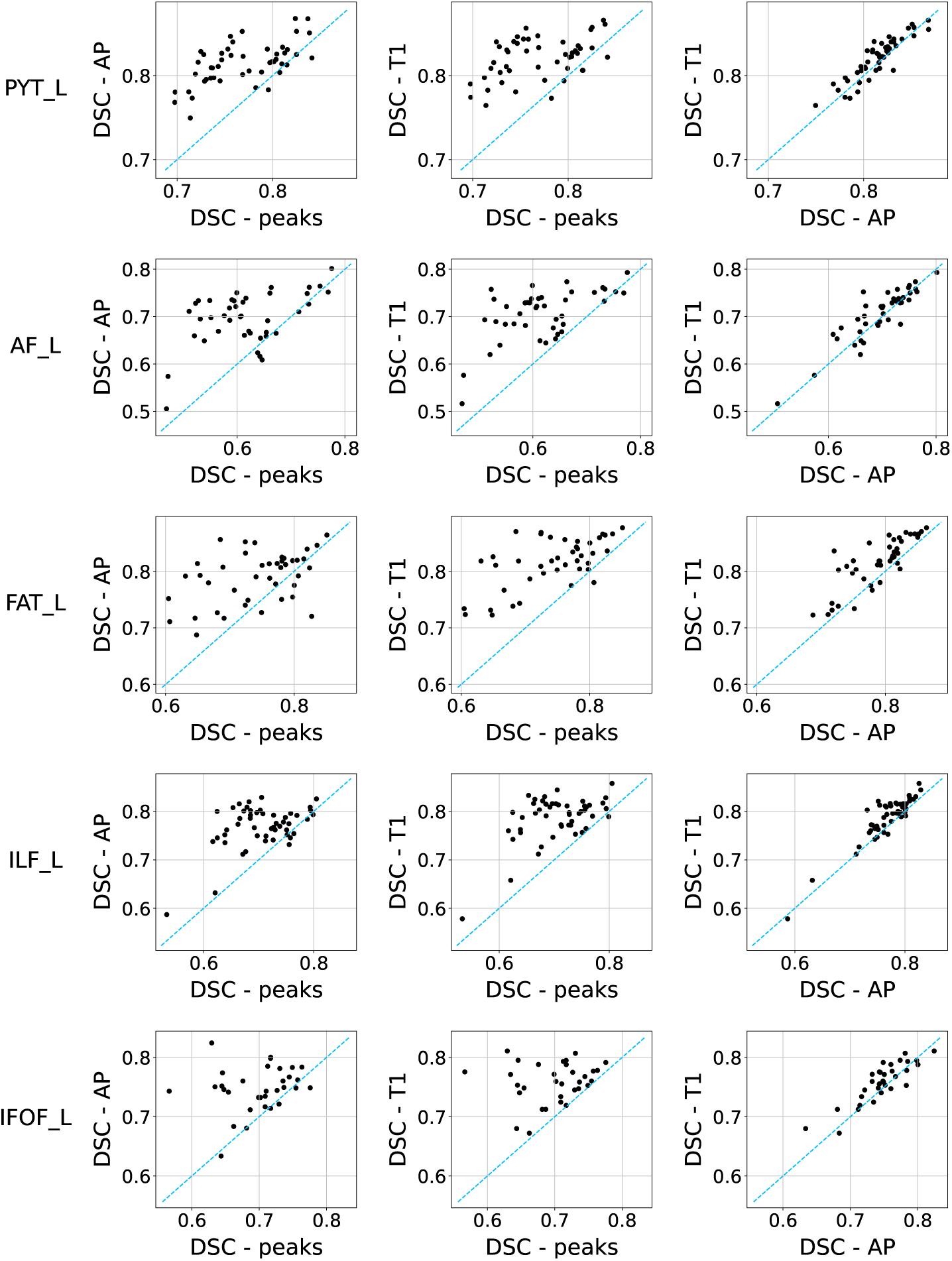
Analysis of input sources. The scatter plots report the relation of DSC measures between the ground truth and the predicted bundle segmentation of models trained with different inputs, where both training and inference are performed on healthy individuals. Each point represents the inference on a single individual. The points located above the diagonal indicate that the input on the y-axis overcomes the performance of the input on the x-axis. Each row refers to a bundle: the pyramidal tract (*PYT_L*), the arcuate fasciculus (*AF_L*), the frontal aslant tract (*FAT_L*), the inferior longitudinal fasciculus (*ILF_L*), and the inferior fronto-occipital fasciculus (*IFOF_L*).

We repeated the analyses for the M-retrain and M-transfer models to understand the influence of different input modalities on segmentation accuracy when applied to glioma patients. We found no significant difference reported between the performances of models trained with different inputs. For this reason, the following analyses only report the results concerning models trained with T1 and AP input.

### 5.4. Domain shift and transfer learning

DSC results are reported graphically in Figure 4 for all input sources and in detail only for the models trained on T1 in Table 2. DSC summary report for all bundles, inputs, and trained models are reported in the supplementary material in Table S1.

**Figure 4:**
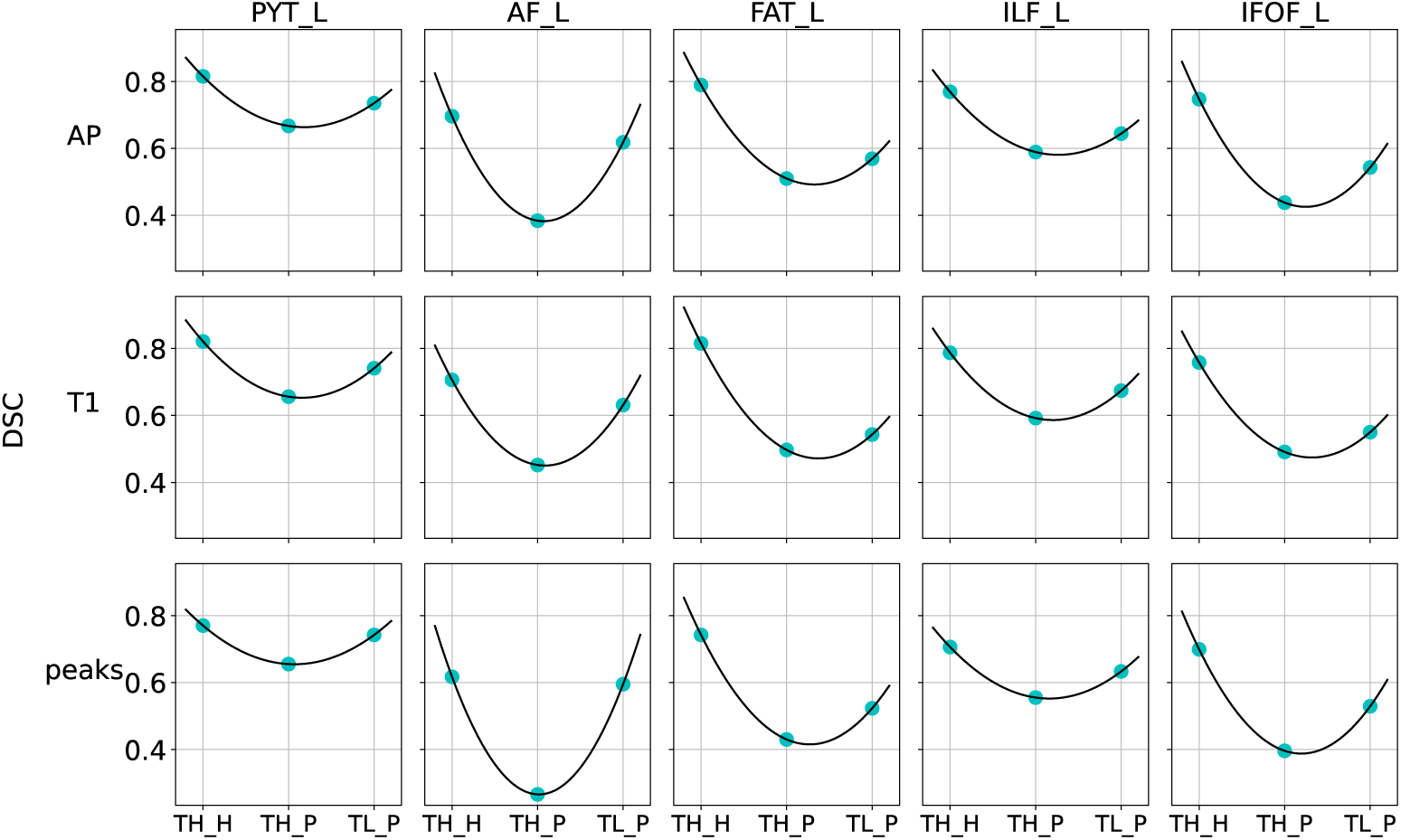
Learning strategies evaluation. On the y-axis: average DSC between ground truth and predictions when models are trained with different inputs. On the x-axis: the different training strategies applied to specific sets of data, in particular: the baseline model M-train, trained on the healthy population and applied to the healthy (TH_H) and glioma patients (TH_P) populations. M-transfer model applied on glioma patients (TL_P). Each column refers to a bundle: the pyramidal tract (*PYT_L*), the arcuate fasciculus (*AF_L*), the frontal aslant tract (*FAT_L*), the inferior longitudinal fasciculus (*ILF_L*), and the inferior fronto-occipital fasciculus (*IFOF_L*).

Figure 4 shows the values of DSC when M-train was used for inference first on the healthy population and then on glioma patients. The third point in the plot refers to the DSC values when inference on glioma patients was carried out with the M-transfer model. The values of DSC for M-retrain predictions on patients are not reported since they are very similar to those obtained with M-transfer.

As expected, the best performance for bundle segmentation was achieved with the M-train model applied to the healthy population. The performance dropped significantly when the same model was used for inference on glioma patients, while M-transfer was effective in partially recovering such a drop. The fine tuning of the original model successfully manages the domain shift, even though the segmentation accuracies on glioma patients were not comparable to those obtained by M-train on healthy individuals. The significance of this recovery in performance was verified with a one-sided Wilcoxon signed-rank test to compare the paired distributions of DSC in the glioma patients’ predictions for M-train and M-transfer. For all the bundles and all the input modalities, we found that M-transfer performs better than M-train in glioma patients, with a *p*-value < 0.01 for all cases, except for the model trained on T1w to segment FAT-L, *p*-value < 0.05, as reported in supplementary materials in Table S3.

The results for M-retrain on glioma patients were analogous to those obtained with the M-transfer model: M-retrain performed better than M-train in patients (verified with a one-sided Wilcoxon signed-rank test reported in supplementary materials, Table S4). However, as reported in Table 2, and as verified by a one-sided Wilcoxon signed-rank test (*p*.value <0.01, supplementary materials, Table S5), M-transfer performed better than M-retrain in the healthy population, better preserving the balance between the two datasets.

5.5. *The recovery of the systematic domain shift*

We studied the relationship between the performance improvement obtained with M-transfer and M-retrain in glioma patients and the spatial domain shift between the two datasets. In particular, we focused on what we labeled systematic domain shift, caused by factors acting homogeneously across the datasets (e.g. as different scanners, acquisition parameters, and segmentation protocols).

We hypothesized that the performance improvement of M-transfer and M-retrain, compared to the baseline model M-train, would be greater in regions exhibiting systematic domain shifts. In particular, this would be true in areas where the domain shift between healthy individuals and patients was more consistently distributed across subjects.

To test this hypothesis, we compared paired voxel maps defined MNI space. The first voxel map, measuring the systematic domain shift, was computed as the absolute difference between the probability maps of the bundles of healthy and clinical datasets (Figures 2 and 5). The second voxel map measured the performance gain provided by the domain adaptation techniques and was defined by two alternative Improvement maps, depending on the training strategy considered. The two different improvement maps were computed to compare the prediction accuracy of the baseline model M-train and of M-retrain and M-transfer, respectively. These were thresholded by selecting values greater than 0 so that only regions where M-retrain and M-transfer improving over M-train were considered. After downsampling the voxel maps to 2mm, we computed a linear regression between the paired voxel maps: the absolute difference of probability maps (the independent variable, or x) versus Improvement map( the dependent variable, or y) for each bundle, for each input modality, and for both transfer learning and retraining. We rejected the null hypothesis of a slope equal to zero with a p-value < 0.01 in all cases, using a one sided T-test with a t-distribution of the test statistic, and considering as alternative hypothesis a slope greater than zero. We also computed the Spearman’s rho correlation coefficient between the absolute difference of probability maps and the Improvement maps that ranged for each bundle and input from 0.40 to 0.60 with p-values of a two-sided T-test all < 0.01. Details about the linear regressions computed, Spearman coefficients, and test statistics are reported in Supplementary materials in Table S6. Taken together, we showed there is a positive relationship between the systematic domain shift and the gain in performance of M-transfer and M-retrain. Figure 5 reports the absolute difference between the probability maps of the two datasets, the improvement map of M-transfer with respect to M-train when they were applied to clinical datasets and the scatter plots between the two.

**Figure 5:**
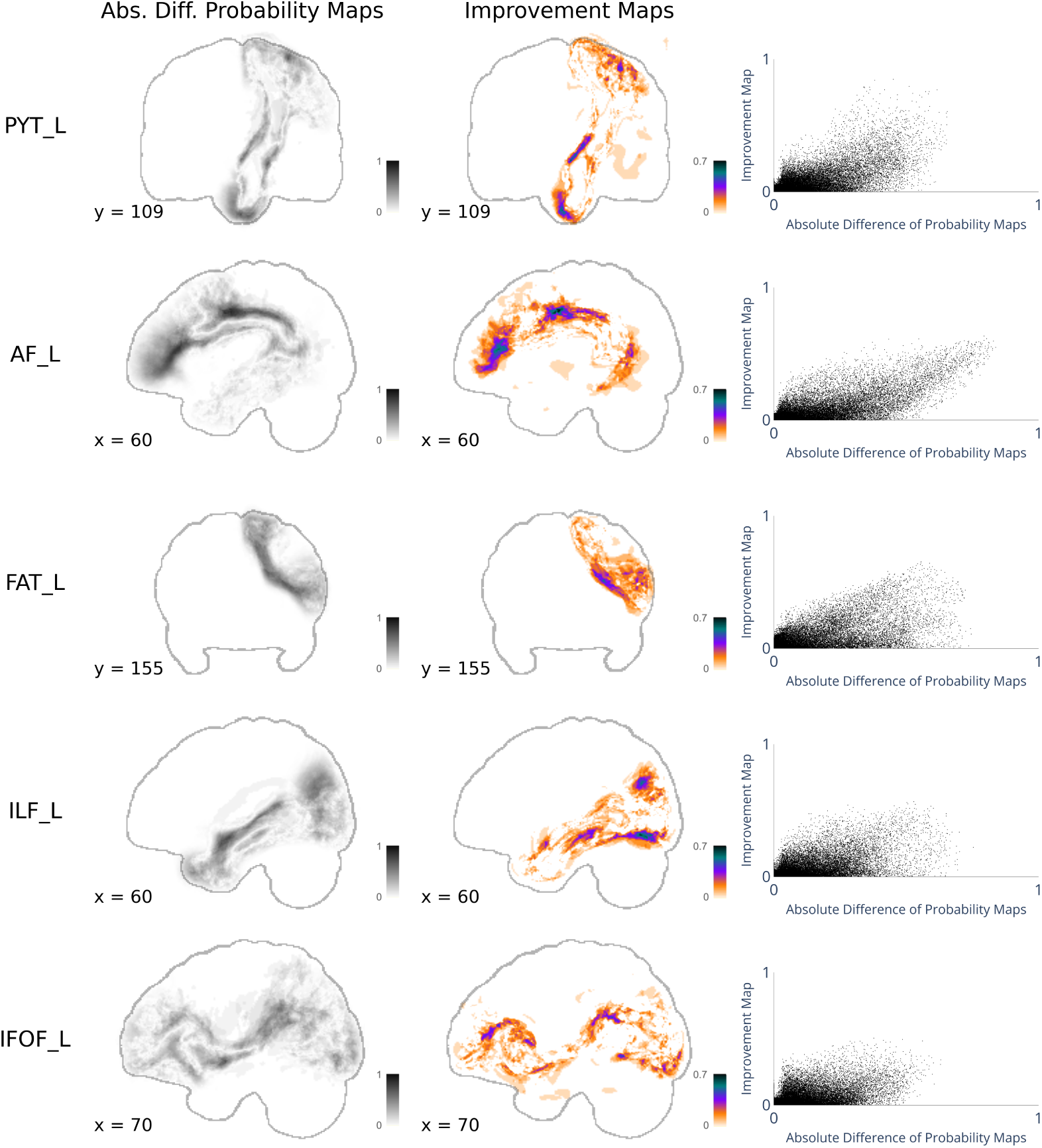
Domain shift and M-transfer Improvement. In the 1st column: the absolute difference between the bundle probability maps of the healthy population and glioma patient datasets. In the 2nd column: the Improvement map, averaged across subjects, and thresholded for values >0, representing the improvement in terms of true positive and negative of M-transfer compared to M-train when they are applied to glioma patients’ set. The considered models are trained with anisotropic power (AP) images. In the 3rd column: the scatterplot represents voxel-wise the relation between these two quantities. Each row refers to a bundle: the pyramidal tract (*PYT_L*), the arcuate fasciculus (*AF_L*), the frontal aslant tract (*FAT_L*), the inferior longitudinal fasciculus (*ILF_L*), and the inferior fronto-occipital fasciculus (*IFOF_L*).

### 5.6. Domain shift and tumor deformation

We investigated whether the domain adaptation techniques M-transfer and M-retrain could recover the domain shift due to tumor-induced deformations. For this analysis, we considered the Warp V-Norm profiles in the healthy subjects and glioma patient datasets. In particular, we inspected the profiles in the glioma patient datasets and we isolated bundles with highly deviant topography (second column in Figure 6). We then verified that the same subjects whose bundle topography was highly deformed (as per Warp V-Norm profiles) had a very low performance in DSC for both the M-transfer and M-retrain’s predictions, and for all data modalities, with a DSC positioned in the lower part of DSC distribution for patients. This was true for all patients with highly deviant bundle topography (Figure 6), suggesting that large alterations in the spatial organization of the bundles were not successfully managed by transfer learning and retraining.

**Figure 6:**
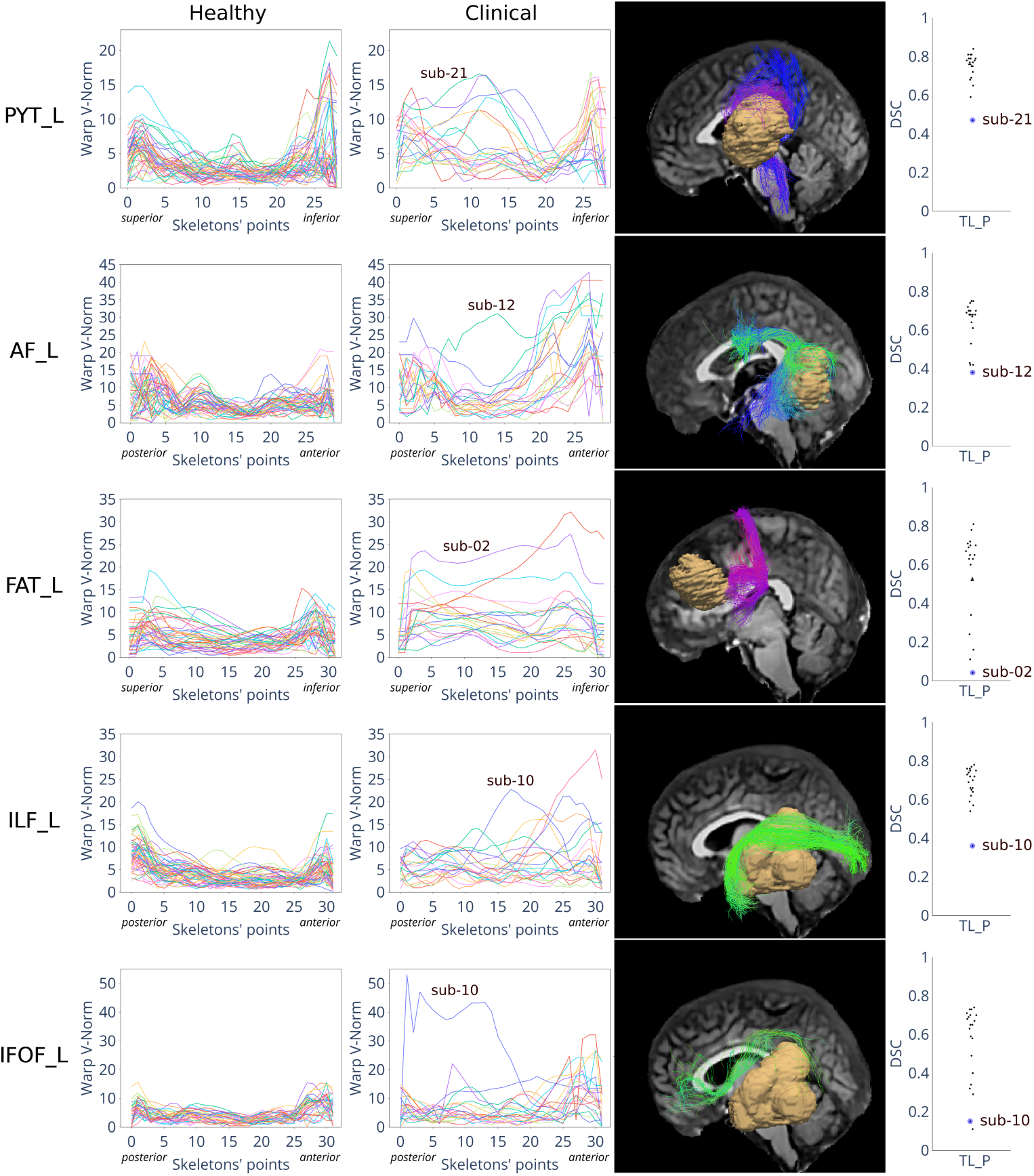
Tumors’ deformations hinder segmentation performances. 1st and 2nd column: Warp V-Norm profiles for the healthy test set and glioma patients. A bundle whose topography is highly deformed is underlined in the clinical Warp-Vnorm profiles, and its bundle-tumor spatial relation is reported in the 3rd column; 4th column: DSCs between ground truth and segmentations predicted by M-transfer trained on T1w and applied to the clinical dataset; The same subjects identified as highly deformed (as per Warp V-Norm profiles) have a very low performance, with a DSC positioned in the lower part of DSC distribution for patients. Each row refers to a bundle, from up to down: the pyramidal tract (*PYT_L*), the arcuate fasciculus (*AF_L*), the frontal aslant tract (*FAT_L*), the inferior longitudinal fasciculus (*ILF_L*), and inferior fronto-occipital fasciculus (*IFOF_L*).

## 6. Discussion

We conducted an empirical investigation to evaluate the efficacy of translating a deep learning bundle segmentation model from healthy subjects to glioma patients. Our findings demonstrate that models trained exclusively on healthy populations exhibit suboptimal performance when applied to a clinical population. Both transfer learning and retraining methodologies proved effective in partially mitigating the domain shift. Notably, transfer learning demonstrated superior performance retention on the healthy dataset compared to retraining.

We observed a substantial decline in the efficacy of bundle segmentation learning models when they were trained on data from healthy subjects and subsequently applied to glioma patients. This performance degradation is consistently observed across various input modalities and white matter bundles examined, as can be observed by Figure 4 and by Table S1 in Supplementary materials. Such findings underscore (i) the necessity for caution when considering the application of deep learning models developed for healthy populations to glioma patients, a strategy widely adopted in the field (Richards et al., 2021; Prasse et al., 2023; Shams et al., 2023); (ii) the necessity of characterizing domain shift.

The first contribution of our work consists in addressing this latter point. We developed a quantitative mapping procedure and the possibility of visualizing how different factors contributed to the observed performance drop. Specifically, we proposed dedicated metrics and visualizations enabling the distinction between systematic (e.g. data preprocessing, acquisition parameters) and neuro-biologically factors (tumor localization ) in shaping the observed domain shift. Importantly, the proposed metrics work both at the single subject level - highlighting patient specific deviations from the normative model - as well as after aggregation, highlighting shifts in the data distribution that are consistent across the whole target datasets. Indeed, we showed that performance improvement was greater in brain regions that were consistently different across datasets. We measured the systemic shift with the difference in probability maps between healthy and clinical datasets bundles, as reported by Figures 2 and 5. This metric aggregates the difference in spatial distribution of all bundles, and measures where domain shift is more consistent between the two datasets. We measured domain shift also at a single subject level with the Warp V-Norm profile that enabled the identification of bundles highly deformed by tumors, as shown in Figure 6.

The second contribution of the present work consists in providing the first empirical investigation evaluating the efficacy of transfer learning for bundle segmentation in glioma patients. Our results suggest that transfer learning may be an effective strategy to manage domain shift, as corroborated by statistically significant recovery of performance observed for all white matter tracts and imaging modalities tested. Intriguingly, a small set of examples in the target clinical domain was enough to trigger the transfer learning mechanism. While the low sample size for the target clinical population may be seen as a limitation, here we argue that it actually represents a strength of our work. First, the sample size adopted in the current study is in line with real-world scenarios. Second, the comparison with state-of-the-art approaches highlights both efficiency and potential of transfer learning. For the pyramidal tract - the only bundle in common with (Moshe et al., 2022)-we observed a mean DSC for M-transfer model of 0.74. Moshe and colleagues (Moshe et al., 2022) reported a mean DSC of 0.64 in the test set, which corresponds to a 15.6% performance decrease using however more than 600 patients, i.e. a 30-fold increase in sample size.

Even though our empirical investigation highlighted the potential of transfer learning, several limitations must be mentioned. First, transfer learning was able to address domain shift, but it did so only partially. As reported above, we observed a performance improvement in brain regions characterized by consistent differences across datasets (Figure 5 ), while patient-specific and tumor-induced bundle deformations were not properly modeled (Figure 6 ). Whether this component of the domain shift can be recovered by increasing sample size in the target domain - or more complex models are needed - remains unclear. Similarly, explaining why more complex imaging modalities underperformed compared to T1w anatomical images requires further investigations. We speculate that the low sample size may act as bottleneck given that the sharp increase in dimensionality was not mirrored by an increase in sample size.

Another limitation of our study is having considered only five bundles. Extending this analysis to a higher number of bundles would make the conclusions even more comprehensive and general. Nevertheless, we underline that the consistency of results between the five bundles tested is already very promising.

Finally, we also observed that M-retrain is as effective as M-transfer in segmenting the bundle in glioma patients. However, it performs worse than M-transfer when applied to the healthy dataset, suggesting that retraining the model’s decoder only is more robust than retraining the whole network.

## 7. Conclusions

Taken together, our results provide additional evidence suggesting that models trained on the healthy population are sub-optimal when applied in clinical settings. Importantly, while we showed that transfer learning and retraining methods are both effective in partially solving domain shift, our results highlight that simple domain adaptation techniques may not be enough, underscoring the importance of a thorough problem characterization.

## 8. CRediT authorship contribution statement

Chiara Riccardi: Conceptualization, Methodology, Software, Visualization, Investigation, Writing – original draft, Writing – review and editing. Ludovico Coletta: Supervision, Writing – original draft, Writing – review and editing. Sofia Ghezzi: Conceptualization, Methodology, Software, Investigation. Gabriele Amorosino: Software, Methodology, Writing – review and editing. Luca Zigiotto: Data curation. Jorge Jovicich: Funding acquisition, Writing – review and editing. Silvio Sarubbo: Data curation. Funding acquisition. Paolo Avesani: Conceptualization, Methodology, Supervision, Project administration, Writing – original draft, Writing – review and editing, Funding acquisition.

## 10. Acknowledgments

This work was partially supported by the grant PAT Reg. n. 764/2021 NeuSurPlan. We also acknowledge the support of the PNRR project FAIR - Future AI Research (PE00000013), under the NRRP MUR program funded by the NextGenerationEU.

## 11. Data availability

The authors do not have permission to share data.

## 12. Compliance with ethical standards

This study was conducted following the ethical standards of the Declaration of Helsinki and was approved by the local ethical committee (authorization ID A734).

## Footnotes

1 https://doi.org/10.18112/openneuro.ds003900.v1.1.1

2 http://trackvis.or^g^

3 https://github.com/scilus/scilpy

4 https://scipy.org/

