## Supplementary materials for "Supervised white matter bundle segmentation in glioma patients with transfer learning"

### 1. Supplementary material

We provide Table S1 reporting the Dice Score Coefficient (DSC) measuring similarity between predicted bundle segmentation and ground truth of models' inferences on the healthy and in glioma patients data. Table S1 reports the average and standard deviation of the DSC for all considered models, M-train, M-transfer, and M-retrain, trained on each considered input, and for segmenting each of the studied bundles. The tested inputs are T1w, anisotropic power, and the three main peaks extracted from Constrained Spherical Deconvolution. The white matter bundles considered are the pyramidal tract (PYT_L), the arcuate fasciculus (AF_L), the frontal aslant tract (FAT_L), the inferior fronto-occipital fasciculus (IFOF_L), and the inferior longitudinal fasciculus (ILF_L).

We also provide tables reporting the statistical tests performed in the paper. Table S2 reports the Wilcoxon signed-rank test performed to compare the DSC performances of models trained with different input modalities and applied to the same test set. We considered the models trained on healthy (M-train) and applied to healthy. Tables S3 and S4 report the Wilcoxon signed-rank test performed to assess that, in glioma patients, transfer learning strategies improved the performances, compared to the model trained only on the healthy population. In particular, the comparison between M-train and M-retrain is reported in Table S4, while Table S3 reports the comparison between M-transfer and M-train in glioma patients. Lastly, Table S5 reports the Wilcoxon signed-rank test to verify that M-transfer performed better than M-retrain in the healthy population. In all Wilcoxon signed-rank tests reported in Table S2, S4, S3, and S5 the paired differences of DSC performances of the models are computed. The tests are one-sided and test the alternative hypothesis that the distribution underlying the differences is stochastically greater than a distribution symmetric about zero. Tables S2, S4, S3, and S5 report details about the tests' results, including the pair of models considered, the median of the DSC of each model, the test statistics, and the $p$.values.

Figure S1 shows the Improvement Maps to compare spatially the performance of model M-train, i.e., trained only on the healthy population, when applied on the healthy test set, across the different input modalities used for the training. The pairwise comparisons across modalities considered were: T1 versus peaks, AP versus peaks, and T1 versus AP. The improvement maps were thresholded for values $> 0$. Darker colors represent where the model trained with the first input modality outperforms the model trained with the second input modality.

Table S6 reports the linear regression performed to assess if there is a positive relationship between the homogeneity of domain shift and the performance improvement of transfer learning strategies, compared to the baseline model trained on healthy, when the models are applied in glioma patients. The linear regression is computed between 1) the absolute difference between the Healthy and Clinical datasets probabilities maps (x, independent variable ), quantifying where the domain shift is homogeneous; and the Improvement Map (y, dependent variable) to compare the performance in glioma patients either between M-retrain and M-train, or between M-transfer and M-train, thresholded for values greater than zero. Table S6 reports details about the linear regression computed, including the slope and the intercept. We report the one-sided T-test that considers as null hypothesis a slope equal to zero and as the alternative hypothesis a slope greater than zero. T-statistics and the p.value of the test are also reported in Table S6. Lastly, in Table S6 we also computed and reported the Spearman coefficients and the related two-sided T-test to compute the p.value representing the probability that two uncorrelated variables produce a dataset with a Spearman correlation at least as extreme as the one computed.

| Model | Modality | Inference | PYT_L | AF_L | FAT_L | ILF_L | IFOF_L |
|---|---|---|---|---|---|---|---|
| M-train | T1w | Healthy | 0.82,0.02 | 0.71, 0.05 | 0.82, 0.04 | 0.79, 0.05 | 0.76, 0.03 |
| M-train | T1w | Patients | 0.66, 0.06 | 0.45,0.11 | 0.50, 0.15 | 0.59,0.09 | 0.49,0.17, |
| M-transfer | T1w | Healthy | 0.75, 0.02 | 0.54, 0.06 | 0.67, 0.07 | 0.67, 0.04 | 0.67, 0.04 |
| M-transfer | T1w | Patients | 0.74, 0.08 | 0.63, 0.12 | 0.54,0.22 | 0.67,0.10 | 0.55, 0.20 |
| M-retrain | T1w | Healthy | 0.67, 0.03 | 0.46, 0.07 | 0.55, 0.08 | 0.59, 0.05 | 0.61, 0.04 |
| M-retrain | T1w | Patients | 0.74,0.09 | 0.65, 0.09 | 0.56, 0.24 | 0.68, 0.1 | 0.56, 0.2 |
| M-train | AP | Healthy | 0.82, 0.02 | 0.7, 0.06 | 0.79, 0.05 | 0.77, 0.04 | 0.75, 0.04 |
| M-train | AP | Patients | 0.67,0.08 | 0.38, 0.09 | 0.51, 0.16 | 0.59, 0.08 | 0.44, 0.16 |
| M-transfer | AP | Healthy | 0.75, 0.02 | 0.56, 0.06 | 0.63, 0.08 | 0.67, 0.04 | 0.69, 0.04 |
| M-transfer | AP | Patients | 0.74, 0.12 | 0.62, 0.1 | 0.57, 0.2 | 0.64, 0.1 | 0.54, 0.19 |
| M-retrain | AP | Healthy | 0.68, 0.03 | 0.48, 0.07 | 0.58, 0.07 | 0.59, 0.04 | 0.62, 0.05 |
| M-retrain | AP | Patients | 0.75, 0.11 | 0.62, 0.1 | 0.58, 0.2 | 0.65, 0.09 | 0.52, 0.2 |
| M-train | peaks | Healthy | 0.77, 0.04 | 0.62, 0.08 | 0.74, 0.07 | 0.71, 0.06 | 0.7, 0.05 |
| M-train | peaks | Patients | 0.66, 0.05 | 0.27, 0.13 | 0.43, 0.17 | 0.56, 0.09 | 0.4, 0.17 |
| M-transfer | peaks | Healthy | 0.72, 0.04 | 0.59, 0.07 | 0.69, 0.06 | 0.62, 0.06 | 0.61, 0.04 |
| M-transfer | peaks | Patients | 0.74, 0.08 | 0.6, 0.1 | 0.52, 0.18 | 0.63, 0.1 | 0.53 0.19 |
| M-retrain | peaks | Healthy | 0.64, 0.05 | 0.44, 0.06 | 0.48, 0.08 | 0.50, 0.06 | 0.55, 0.03 |
| M-retrain | peaks | Patients | 0.75, 0.09 | 0.63, 0.1 | 0.56, 0.17 | 0.66, 0.1 | 0.56, 0.2 |

**Table S1. DSC metrics: Mean and Standard deviation of DSC between ground truth and predicted bundle segmentations of models M-train, M-transfer, and M-retrain trained by using as input images of T1w, AP, or peaks. The models are applied to the Healthy population (Healthy) or glioma patient datasets (Patients). The bundles considered are the pyramidal tract (PYT_L), the arcuate fasciculus (AF_L), the frontal aslant tract (FAT_L), the inferior fronto-occipital fasciculus (IFOF_L), and the inferior longitudinal fasciculus (ILF_L).**

| Model | Inference | Bundle | Paired Differences | Medians 1st Input | Medians 2nd Input | test statistics | $p$.value |
|---|---|---|---|---|---|---|---|
| M-train | Healthy | PYT_L | $d_i = DSC_{T1w_i} - DSC_{peaks_i}$ | $Med(DSC_{T1w}) = 0.83$ | $Med(DSC_{peaks}) = 0.77$ | 1151 | 3.86E-09 |
| M-train | Healthy | AF_L | $d_i = DSC_{T1w_i} - DSC_{peaks_i}$ | $Med(DSC_{T1w}) = 0.72$ | $Med(DSC_{peaks}) = 0.61$ | 893 | 1.69E-08 |
| M-train | Healthy | FAT_L | $d_i = DSC_{T1w_i} - DSC_{peaks_i}$ | $Med(DSC_{T1w}) = 0.82$ | $Med(DSC_{peaks}) = 0.76$ | 896 | 1.37E-08 |
| M-train | Healthy | ILF_L | $d_i = DSC_{T1w_i} - DSC_{peaks_i}$ | $Med(DSC_{T1w}) = 0.80$ | $Med(DSC_{peaks}) = 0.71$ | 1272 | 4.53E-10 |
| M-train | Healthy | IFOF_L | $d_i = DSC_{T1w_i} - DSC_{peaks_i}$ | $Med(DSC_{T1w}) = 0.76$ | $Med(DSC_{peaks}) = 0.71$ | 528 | 3.98E-07 |
| M-train | Healthy | PYT_L | $d_i = DSC_{AP_i} - DSC_{peaks_i}$ | $Med(DSC_{AP}) = 0.82$ | $Med(DSC_{peaks}) = 0.77$ | 1148 | 4.63E-09 |
| M-train | Healthy | AF_L | $d_i = DSC_{AP_i} - DSC_{peaks_i}$ | $Med(DSC_{AP}) = 0.70$ | $Med(DSC_{peaks}) = 0.61$ | 850 | 3.13E-07 |
| M-train | Healthy | FAT_L | $d_i = DSC_{AP_i} - DSC_{peaks_i}$ | $Med(DSC_{AP}) = 0.80$ | $Med(DSC_{peaks}) = 0.76$ | 791 | 1.09E-05 |
| M-train | Healthy | ILF_L | $d_i = DSC_{AP_i} - DSC_{peaks_i}$ | $Med(DSC_{AP}) = 0.77$ | $Med(DSC_{peaks}) = 0.71$ | 1242 | 2.68E-09 |
| M-train | Healthy | IFOF_L | $d_i = DSC_{AP_i} - DSC_{peaks_i}$ | $Med(DSC_{AP}) = 0.75$ | $Med(DSC_{peaks}) = 0.71$ | 492 | 1.01E-05 |
| M-train | Healthy | PYT_L | $d_i = DSC_{T1w_i} - DSC_{AP_i}$ | $Med(DSC_{T1w}) = 0.83$ | $Med(DSC_{AP}) = 0.82$ | 922 | 3.07E-04 |
| M-train | Healthy | AF_L | $d_i = DSC_{T1w_i} - DSC_{AP_i}$ | $Med(DSC_{T1w}) = 0.72$ | $Med(DSC_{AP}) = 0.70$ | 624 | 1.55E-02 |
| M-train | Healthy | FAT_L | $d_i = DSC_{T1w_i} - DSC_{AP_i}$ | $Med(DSC_{T1w}) = 0.82$ | $Med(DSC_{AP}) = 0.80$ | 846 | 4.06E-07 |
| M-train | Healthy | ILF_L | $d_i = DSC_{T1w_i} - DSC_{AP_i}$ | $Med(DSC_{T1w}) = 0.80$ | $Med(DSC_{AP}) = 0.77$ | 1186 | 5.96E-08 |
| M-train | Healthy | IFOF_L | $d_i = DSC_{T1w_i} - DSC_{AP_i}$ | $Med(DSC_{T1w}) = 0.76$ | $Med(DSC_{AP}) = 0.75$ | 414 | 2.52E-03 |

**Table S2. Wilcoxon signed-rank test to compare input data modalities. The test compared the DSC performance of models trained with different input modalities, considering the case of training and inference on the healthy population. The test considered the paired differences of DSC performances of models using different input modalities and applied to the same test set. The test was one-sided and tested the alternative hypothesis that the distribution underlying the differences was stochastically greater than a distribution symmetric about zero. The test was performed with Scipy (`https://scipy.org/`). In the table are reported: the bundle; the considered paired differences of the DSC performances of models using different input modalities; the median values of DSC performances of the models compared; the test statistics that refers to the sum of the ranks of the differences above zero; the $p$. value. The bundles considered are the pyramidal tract (PYT_L), the arcuate fasciculus (AF_L), the frontal aslant tract (FAT_L), the inferior fronto-occipital fasciculus (IFOF_L), and the inferior longitudinal fasciculus (ILF_L).**

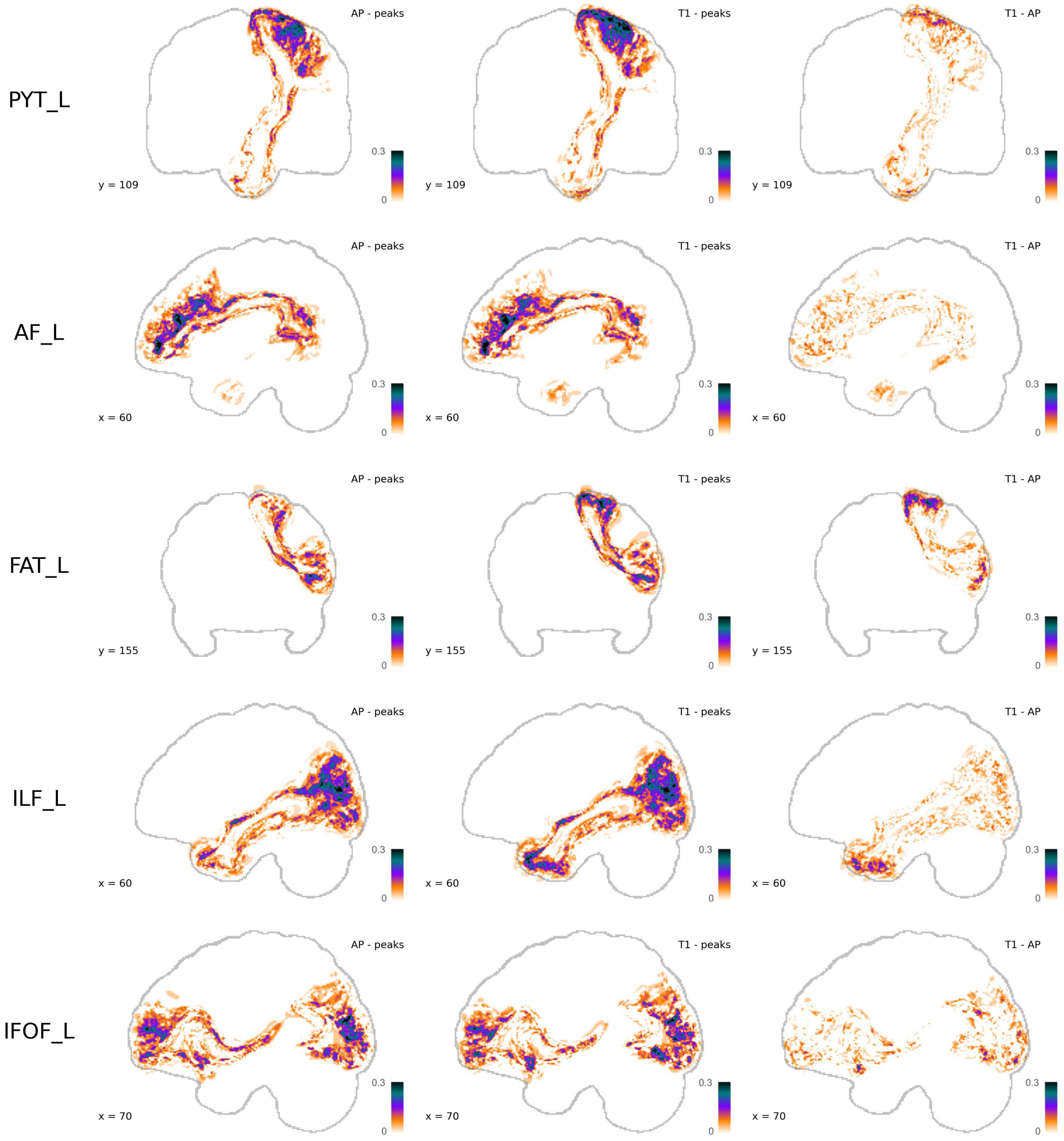


**Fig. S1. Comparison across input modalities with Improvement maps. Improvement maps showing the performance comparison across models trained with different input images. We considered M-train models, trained on the healthy dataset when they are applied to the healthy test set. The improvement maps are thresholded to have values greater than 0. Darker colors represent where: 1) the AP model outperforms the peak model (first column); 2) the T1 model outperforms the peak model (second column); 3) the T1 model outperforms the AP model (third column). Each row refers to a bundle, from up to down: the pyramidal tract (PYT_L), the arcuate fasciculus (AF_L), the frontal aslant tract (FAT_L), the inferior longitudinal fasciculus (ILF_L), and the inferior fronto-occipital fasciculus (IFOF_L).**

| Modality | Inference | Bundle | Paired Differences | Medians 1st Model | Medians 2nd Model | test statistics | $p$.value |
|---|---|---|---|---|---|---|---|
| T1w | Patients | PYT_L | $d_i = DSC_{TL_i} - DSC_{TH_i}$ | $Med(DSC_{TL}) = 0.77$ | $Med(DSC_{TH}) = 0.67$ | 227 | 3.34E-06 |
| T1w | Patients | AF_L | $d_i = DSC_{TL_i} - DSC_{TH_i}$ | $Med(DSC_{TL}) = 0.68$ | $Med(DSC_{TH}) = 0.48$ | 231 | 4.77E-07 |
| T1w | Patients | FAT_L | $d_i = DSC_{TL_i} - DSC_{TH_i}$ | $Med(DSC_{TL}) = 0.63$ | $Med(DSC_{TH}) = 0.55$ | 166 | 4.11E-02 |
| T1w | Patients | ILF_L | $d_i = DSC_{TL_i} - DSC_{TH_i}$ | $Med(DSC_{TL}) = 0.7$ | $Med(DSC_{TH}) = 0.60$ | 228 | 2.38E-06 |
| T1w | Patients | IFOF_L | $d_i = DSC_{TL_i} - DSC_{TH_i}$ | $Med(DSC_{TL}) = 0.65$ | $Med(DSC_{TH}) = 0.57$ | 224 | 9.06E-06 |
| AP | Patients | PYT_L | $d_i = DSC_{TL_i} - DSC_{TH_i}$ | $Med(DSC_{TL}) = 0.78$ | $Med(DSC_{TH}) = 0.68$ | 201 | 3.15E-05 |
| AP | Patients | AF_L | $d_i = DSC_{TL_i} - DSC_{TH_i}$ | $Med(DSC_{TL}) = 0.64$ | $Med(DSC_{TH}) = 0.38$ | 210 | 9.54E-07 |
| AP | Patients | FAT_L | $d_i = DSC_{TL_i} - DSC_{TH_i}$ | $Med(DSC_{TL}) = 0.63$ | $Med(DSC_{TH}) = 0.55$ | 173 | 4.72E-03 |
| AP | Patients | ILF_L | $d_i = DSC_{TL_i} - DSC_{TH_i}$ | $Med(DSC_{TL}) = 0.68$ | $Med(DSC_{TH}) = 0.6$ | 185 | 8.45E-04 |
| AP | Patients | IFOF_L | $d_i = DSC_{TL_i} - DSC_{TH_i}$ | $Med(DSC_{TL}) = 0.63$ | $Med(DSC_{TH}) = 0.5$ | 209 | 1.91E-06 |
| peaks | Patients | PYT_L | $d_i = DSC_{TL_i} - DSC_{TH_i}$ | $Med(DSC_{TL}) = 0.77$ | $Med(DSC_{TH}) = 0.66$ | 206 | 6.68E-06 |
| peaks | Patients | AF_L | $d_i = DSC_{TL_i} - DSC_{TH_i}$ | $Med(DSC_{TL}) = 0.64$ | $Med(DSC_{TH}) = 0.27$ | 210 | 9.54E-07 |
| peaks | Patients | FAT_L | $d_i = DSC_{TL_i} - DSC_{TH_i}$ | $Med(DSC_{TL}) = 0.56$ | $Med(DSC_{TH}) = 0.46$ | 192 | 2.41E-04 |
| peaks | Patients | ILF_L | $d_i = DSC_{TL_i} - DSC_{TH_i}$ | $Med(DSC_{TL}) = 0.66$ | $Med(DSC_{TH}) = 0.56$ | 202 | 2.38E-05 |
| peaks | Patients | IFOF_L | $d_i = DSC_{TL_i} - DSC_{TH_i}$ | $Med(DSC_{TL}) = 0.60$ | $Med(DSC_{TH}) = 0.47$ | 209 | 1.91E-06 |

**Table S3. Wilcoxon signed-rank test to compare M-transfer and M-train performances in patients. The test compared the DSC performance of models M-train and M-transfer, both trained for the segmentation of the same bundle, with the same input modality, when they were applied to the same set of glioma patients. Models M-train and M-transfer are indicated in the table respectively by TH and TL. The test considered the paired differences in DSC performances between these two models. The test was one-sided and tested the alternative hypothesis that the distribution underlying the differences was stochastically greater than a distribution symmetric about zero. The test was performed with Scipy (`https://scipy.org/`). In the table are reported: the input modality; the inference set; the bundle; the considered paired differences of the DSC performances between different models; the median values of DSC performances of the models compared; the test statistic that refers to the sum of the ranks of the differences above zero; the $p$. value. The input modalities are T1w, anisotropic power (AP), and peaks. The bundles considered are the pyramidal tract (PYT_L), the arcuate fasciculus (AF_L), the frontal aslant tract (FAT_L), the inferior fronto-occipital fasciculus (IFOF_L), and the inferior longitudinal fasciculus (ILF_L).**

| Modality | Inference | Bundle | Paired Differences | Medians 1st Model | Medians 2nd Model | test statistics | $p$.value |
|---|---|---|---|---|---|---|---|
| T1w | Patients | PYT_L | $d_i = DSC_{RT_i} - DSC_{TH_i}$ | $Med(DSC_{RT}) = 0.77$ | $Med(DSC_{TH}) = 0.67$ | 227 | 3.34E-06 |
| T1w | Patients | AF_L | $d_i = DSC_{RT_i} - DSC_{TH_i}$ | $Med(DSC_{RT}) = 0.68$ | $Med(DSC_{TH}) = 0.48$ | 231 | 4.77E-07 |
| T1w | Patients | FAT_L | $d_i = DSC_{RT_i} - DSC_{TH_i}$ | $Med(DSC_{RT}) = 0.68$ | $Med(DSC_{TH}) = 0.55$ | 170 | 2.98E-02 |
| T1w | Patients | ILF_L | $d_i = DSC_{RT_i} - DSC_{TH_i}$ | $Med(DSC_{RT}) = 0.72$ | $Med(DSC_{TH}) = 0.60$ | 228 | 2.38E-06 |
| T1w | Patients | IFOF_L | $d_i = DSC_{RT_i} - DSC_{TH_i}$ | $Med(DSC_{RT}) = 0.65$ | $Med(DSC_{TH}) = 0.57$ | 219 | 3.34E-05 |
| AP | Patients | PYT_L | $d_i = DSC_{RT_i} - DSC_{TH_i}$ | $Med(DSC_{RT}) = 0.78$ | $Med(DSC_{TH}) = 0.68$ | 208 | 2.86E-06 |
| AP | Patients | AF_L | $d_i = DSC_{RT_i} - DSC_{TH_i}$ | $Med(DSC_{RT}) = 0.66$ | $Med(DSC_{TH}) = 0.38$ | 210 | 9.54E-07 |
| AP | Patients | FAT_L | $d_i = DSC_{RT_i} - DSC_{TH_i}$ | $Med(DSC_{RT}) = 0.66$ | $Med(DSC_{TH}) = 0.55$ | 172 | 5.34E-03 |
| AP | Patients | ILF_L | $d_i = DSC_{RT_i} - DSC_{TH_i}$ | $Med(DSC_{RT}) = 0.66$ | $Med(DSC_{TH}) = 0.60$ | 194 | 1.61E-04 |
| AP | Patients | IFOF_L | $d_i = DSC_{RT_i} - DSC_{TH_i}$ | $Med(DSC_{RT}) = 0.58$ | $Med(DSC_{TH}) = 0.50$ | 168 | 8.59E-03 |
| peaks | Patients | PYT_L | $d_i = DSC_{RT_i} - DSC_{TH_i}$ | $Med(DSC_{RT}) = 0.77$ | $Med(DSC_{TH}) = 0.66$ | 203 | 1.81E-05 |
| peaks | Patients | AF_L | $d_i = DSC_{RT_i} - DSC_{TH_i}$ | $Med(DSC_{RT}) = 0.67$ | $Med(DSC_{TH}) = 0.27$ | 210 | 9.54E-07 |
| peaks | Patients | FAT_L | $d_i = DSC_{RT_i} - DSC_{TH_i}$ | $Med(DSC_{RT}) = 0.57$ | $Med(DSC_{TH}) = 0.46$ | 200 | 4.10E-05 |
| peaks | Patients | ILF_L | $d_i = DSC_{RT_i} - DSC_{TH_i}$ | $Med(DSC_{RT}) = 0.68$ | $Med(DSC_{TH}) = 0.56$ | 207 | 4.77E-06 |
| peaks | Patients | IFOF_L | $d_i = DSC_{RT_i} - DSC_{TH_i}$ | $Med(DSC_{RT}) = 0.64$ | $Med(DSC_{TH}) = 0.47$ | 210 | 9.54E-07 |

**Table S4. Wilcoxon signed-rank Test to compare M-retrain and M-train performances in patients. The test compared the DSC performance of models M-retrain and M-transfer, both trained for the segmentation of the same bundle, with the same input modality, when they were applied to the same set of glioma patients. Models M-train and M-retrain are indicated in the table respectively by TH and RT. The test considered the paired differences in DSC performances between these two models. The test was one-sided and tested the alternative hypothesis that the distribution underlying the differences was stochastically greater than a distribution symmetric about zero. The test was performed with Scipy (`https://scipy.org/`). In the table are reported: the input modality; the inference set; the bundle; the considered paired differences of the DSC performances between different models; the median values of DSC performances of the models compared; the test statistic that refers to the sum of the ranks of the differences above zero; the $p$. value. The input modalities are T1w, anisotropic power (AP), and peaks. The bundles considered are the pyramidal tract (PYT_L), the arcuate fasciculus (AF_L), the frontal aslant tract (FAT_L), the inferior fronto-occipital fasciculus (IFOF_L), and the inferior longitudinal fasciculus (ILF_L).**

| Modality | Inference | Bundle | Paired Differences | Medians 1st Model | Medians 2nd Model | test statistics | $p$.value |
|---|---|---|---|---|---|---|---|
| T1w | Healthy | PYT_L | $d_i = DSC_{TL_i} - DSC_{RT_i}$ | $Med(DSC_{TL}) = 0.75$ | $Med(DSC_{RT}) = 0.67$ | 1176 | 7.84E-10 |
| T1w | Healthy | AF_L | $d_i = DSC_{TL_i} - DSC_{RT_i}$ | $Med(DSC_{TL}) = 0.54$ | $Med(DSC_{RT}) = 0.47$ | 903 | 8.17E-09 |
| T1w | Healthy | FAT_L | $d_i = DSC_{TL_i} - DSC_{RT_i}$ | $Med(DSC_{TL}) = 0.68$ | $Med(DSC_{RT}) = 0.57$ | 902 | 8.80E-09 |
| T1w | Healthy | ILF_L | $d_i = DSC_{TL_i} - DSC_{RT_i}$ | $Med(DSC_{TL}) = 0.67$ | $Med(DSC_{RT}) = 0.59$ | 1275 | 3.71E-10 |
| T1w | Healthy | IFOF_L | $d_i = DSC_{TL_i} - DSC_{RT_i}$ | $Med(DSC_{TL}) = 0.67$ | $Med(DSC_{RT}) = 0.60$ | 525.5 | 4.95E-07 |
| AP | Healthy | PYT_L | $d_i = DSC_{TL_i} - DSC_{RT_i}$ | $Med(DSC_{TL}) = 0.75$ | $Med(DSC_{RT}) = 0.69$ | 1176 | 7.70E-10 |
| AP | Healthy | AF_L | $d_i = DSC_{TL_i} - DSC_{RT_i}$ | $Med(DSC_{TL}) = 0.56$ | $Med(DSC_{RT}) = 0.47$ | 820 | 1.77E-08 |
| AP | Healthy | FAT_L | $d_i = DSC_{TL_i} - DSC_{RT_i}$ | $Med(DSC_{TL}) = 0.64$ | $Med(DSC_{RT}) = 0.60$ | 849 | 3.26E-07 |
| AP | Healthy | ILF_L | $d_i = DSC_{TL_i} - DSC_{RT_i}$ | $Med(DSC_{TL}) = 0.67$ | $Med(DSC_{RT}) = 0.59$ | 1275 | 3.70E-10 |
| AP | Healthy | IFOF_L | $d_i = DSC_{TL_i} - DSC_{RT_i}$ | $Med(DSC_{TL}) = 0.69$ | $Med(DSC_{RT}) = 0.63$ | 528 | 3.83E-07 |
| peaks | Healthy | PYT_L | $d_i = DSC_{TL_i} - DSC_{RT_i}$ | $Med(DSC_{TL}) = 0.71$ | $Med(DSC_{RT}) = 0.65$ | 1176 | 7.83E-10 |
| peaks | Healthy | AF_L | $d_i = DSC_{TL_i} - DSC_{RT_i}$ | $Med(DSC_{TL}) = 0.57$ | $Med(DSC_{RT}) = 0.44$ | 903 | 8.21E-09 |
| peaks | Healthy | FAT_L | $d_i = DSC_{TL_i} - DSC_{RT_i}$ | $Med(DSC_{TL}) = 0.70$ | $Med(DSC_{RT}) = 0.50$ | 902 | 8.76E-09 |
| peaks | Healthy | ILF_L | $d_i = DSC_{TL_i} - DSC_{RT_i}$ | $Med(DSC_{TL}) = 0.62$ | $Med(DSC_{RT}) = 0.5$ | 1275 | 3.73E-10 |
| peaks | Healthy | IFOF_L | $d_i = DSC_{TL_i} - DSC_{RT_i}$ | $Med(DSC_{TL}) = 0.62$ | $Med(DSC_{RT}) = 0.55$ | 492.5 | 8.16E-07 |

**Table S5. Wilcoxon signed-rank test to compare M-retrain and M-transfer performances in the healthy population. The test compared the DSC performance of models M-retrain and M-transfer, both trained for the segmentation of the same bundle, with the same input modality, when they were applied to the same set of healthy population. Models M-retrain and M-transfer are indicated in the table respectively by RT and TL. The test considered the paired differences in DSC performances between these two models. The test was one-sided and tested the alternative hypothesis that the distribution underlying the differences was stochastically greater than a distribution symmetric about zero. The test was performed with Scipy (`https://scipy.org/`). The table reports: the input modality; the inference set; the bundle; the considered paired differences of the DSC performances between different models; the median values of DSC performances of the models compared; the test statistic that refers to the sum of the ranks of the differences above zero; the $p$. value. The input modalities are T1w, anisotropic power (AP), and peaks. The bundles considered are the pyramidal tract (PYT_L), the arcuate fasciculus (AF_L), the frontal aslant tract (FAT_L), the inferior fronto-occipital fasciculus (IFOF_L), and the inferior longitudinal fasciculus (ILF_L).**

| Bundle | Inference | Modality | y | x | slope | intercept | std.err slope | t-statistics slope | p. value slope | Spearman | t-statistics Spearman | p. value Spearman |
|---|---|---|---|---|---|---|---|---|---|---|---|---|
| PYT_L | Patients | T1 | Improvement Map(RT, TH) | Difference in Prob. Maps | 0.622 | 0.021 | 0.006 | 99.64 | 0 | 0.50 | 87.19 | 0 |
| AF_L | Patients | T1 | Improvement Map(RT, TH) | Difference in Prob. Maps | 0.589 | 0.007 | 0.004 | 147.76 | 0 | 0.62 | 122.06 | 0 |
| IFOF_L | Patients | T1 | Improvement Map(RT, TH) | Difference in Prob. Maps | 0.407 | 0.016 | 0.004 | 98.25 | 0 | 0.45 | 79.65 | 0 |
| FAT_L | Patients | T1 | Improvement Map(RT, TH) | Difference in Prob. Maps | 0.496 | 0.026 | 0.004 | 110.94 | 0 | 0.65 | 121.26 | 0 |
| ILF_L | Patients | T1 | Improvement Map(RT, TH) | Difference in Prob. Maps | 0.541 | 0.014 | 0.004 | 132.06 | 0 | 0.64 | 127.25 | 0 |
| PYT_L | Patients | AP | Improvement Map(RT, TH) | Difference in Prob. Maps | 0.575 | 0.013 | 0.004 | 157.21 | 0 | 0.57 | 114.07 | 0 |
| AF_L | Patients | AP | Improvement Map(RT, TH) | Difference in Prob. Maps | 0.475 | 0.008 | 0.003 | 152.35 | 0 | 0.62 | 119.62 | 0 |
| IFOF_L | Patients | AP | Improvement Map(RT, TH) | Difference in Prob. Maps | 0.308 | 0.025 | 0.003 | 99.67 | 0 | 0.42 | 77.80 | 0 |
| FAT_L | Patients | AP | Improvement Map(RT, TH) | Difference in Prob. Maps | 0.417 | 0.03 | 0.004 | 109.76 | 0 | 0.56 | 96.84 | 0 |
| ILF_L | Patients | AP | Improvement Map(RT, TH) | Difference in Prob. Maps | 0.348 | 0.029 | 0.004 | 89.10 | 0 | 0.48 | 79.89 | 0 |
| PYT_L | Patients | peaks | Improvement Map(RT, TH) | Difference in Prob. Maps | 0.66 | 0.011 | 0.005 | 135.19 | 0 | 0.56 | 105.60 | 0 |
| AF_L | Patients | peaks | Improvement Map(RT, TH) | Difference in Prob. Maps | 0.339 | 0.042 | 0.005 | 65.17 | 0 | 0.44 | 61.14 | 0 |
| IFOF_L | Patients | peaks | Improvement Map(RT, TH) | Difference in Prob. Maps | 0.561 | 0.042 | 0.005 | 107.47 | 0 | 0.50 | 109.93 | 0 |
| FAT_L | Patients | peaks | Improvement Map(RT, TH) | Difference in Prob. Maps | 0.217 | 0.033 | 0.003 | 73.92 | 0 | 0.54 | 83.90 | 0 |
| ILF_L | Patients | peaks | Improvement Map(RT, TH) | Difference in Prob. Maps | 0.448 | 0.052 | 0.006 | 77.20 | 0 | 0.50 | 86.83 | 0 |
| PYT_L | Patients | T1 | Improvement Map(TL, TH) | Difference in Prob. Maps | 0.594 | 0.017 | 0.006 | 95.19 | 0 | 0.50 | 86.48 | 0 |
| AF_L | Patients | T1 | Improvement Map(TL, TH) | Difference in Prob. Maps | 0.578 | 0.004 | 0.004 | 139.63 | 0 | 0.61 | 115.67 | 0 |
| IFOF_L | Patients | T1 | Improvement Map(TL, TH) | Difference in Prob. Maps | 0.399 | 0.012 | 0.004 | 93.90 | 0 | 0.43 | 72.58 | 0 |
| FAT_L | Patients | T1 | Improvement Map(TL, TH) | Difference in Prob. Maps | 0.457 | 0.026 | 0.005 | 99.23 | 0 | 0.60 | 104.22 | 0 |
| ILF_L | Patients | T1 | Improvement Map(TL, TH) | Difference in Prob. Maps | 0.491 | 0.015 | 0.004 | 119.73 | 0 | 0.60 | 115.45 | 0 |
| PYT_L | Patients | AP | Improvement Map(TL, TH) | Difference in Prob. Maps | 0.5 | 0.014 | 0.003 | 143.23 | 0 | 0.56 | 109.65 | 0 |
| AF_L | Patients | AP | Improvement Map(TL, TH) | Difference in Prob. Maps | 0.446 | 0.008 | 0.003 | 147.13 | 0 | 0.62 | 114.21 | 0 |
| IFOF_L | Patients | AP | Improvement Map(TL, TH) | Difference in Prob. Maps | 0.305 | 0.025 | 0.004 | 86.64 | 0 | 0.46 | 80.37 | 0 |
| FAT_L | Patients | AP | Improvement Map(TL, TH) | Difference in Prob. Maps | 0.367 | 0.032 | 0.004 | 98.83 | 0 | 0.51 | 84.66 | 0 |
| ILF_L | Patients | AP | Improvement Map(TL, TH) | Difference in Prob. Maps | 0.353 | 0.026 | 0.004 | 88.46 | 0 | 0.46 | 75.02 | 0 |
| PYT_L | Patients | peaks | Improvement Map(TL, TH) | Difference in Prob. Maps | 0.686 | 0.006 | 0.005 | 137.45 | 0 | 0.56 | 104.72 | 0 |
| AF_L | Patients | peaks | Improvement Map(TL, TH) | Difference in Prob. Maps | 0.31 | 0.044 | 0.006 | 55.39 | 0 | 0.41 | 52.91 | 0 |
| IFOF_L | Patients | peaks | Improvement Map(TL, TH) | Difference in Prob. Maps | 0.526 | 0.037 | 0.005 | 100.22 | 0 | 0.47 | 98.28 | 0 |
| FAT_L | Patients | peaks | Improvement Map(TL, TH) | Difference in Prob. Maps | 0.181 | 0.031 | 0.003 | 63.23 | 0 | 0.49 | 70.50 | 0 |
| ILF_L | Patients | peaks | Improvement Map(TL, TH) | Difference in Prob. Maps | 0.408 | 0.049 | 0.006 | 71.87 | 0 | 0.48 | 79.25 | 0 |

**Table S6. Linear Regression between Improvement Maps and Absolute Difference in Probability maps. The Table reports the linear regression performed between the Improvement Map (y, dependent variable) and the absolute difference between the Healthy and Clinical datasets probabilities maps (x, independent variable ). The Improvement Maps compare the performance in glioma patients either between M-retrain and M-train, or between M-transfer and M-train and are thresholded for values greater than zero. The training strategies M-train, M-transfer, and M-retrain are indicated in the table respectively with TH, TL, and RT. The table reports the bundle, the used data modality, which are the dependent and independent variables of the linear regression, specifying which models are compared by each Improvement Map, and information about the linear regressions computed, including the slope, the intercept, the standard error of the slope. We performed also a one-sided T-test considering the null hypothesis of a slope equal to zero and as the alternative hypothesis a slope greater than zero. The t-statistics and the p.value of the test are reported. We also computed and reported the Spearman coefficients and the related two-sided T-test to compute the p.value representing the probability that two uncorrelated variables produce a dataset with a Spearman correlation at least as extreme as the one computed.**
